# FRET monitoring of transcription factor activities in living bacteria

**DOI:** 10.1101/2022.01.14.476424

**Authors:** Pengchao Wang, Guangming Zhang, Zeling Xu, Zhe Chen, Xiaohong Liu, Chenyin Wang, Chaogu Zheng, Jiangyun Wang, Hongmin Zhang, Aixin Yan

## Abstract

Bacteria adapt to the constantly changing environments largely by transcriptional regulation through the activities of various transcription factors (TFs). However, techniques that monitor the in situ TF-promoter interactions in living bacteria are lacking. Herein, we developed a whole-cell TF-promoter binding assay based on the intermolecular Förster resonance energy transfer (FRET) between a fluorescent unnatural amino acid CouA which is genetically encoded into defined sites in TFs and the live cell fluorescent nucleic acid stain SYTO 9. We show that this new FRET pair monitors the intricate TF-promoter interactions elicited by various types of signal transduction systems with specificity and sensitivity. Furthermore, the assay is applicable to identify novel modulators of the regulatory systems of interest and monitor TF activities in bacteria colonized in *C. elegans*. In conclusion, we established a tractable and sensitive TF-promoter binding assay in living bacteria which not only complements currently available approaches for DNA-protein interactions but also provides novel opportunities for functional annotation of bacterial signal transduction systems and studies of the bacteria-host interface.

## Introduction

Pathogenic bacteria encounter diverse environmental and physiological stresses in their natural habitats and during infection of the human hosts, such as oxygen fluctuation, nutrient scarce, hyper-osmolarity, bile salts, bactericidal agents, etc. (1). To survive these hostile environments, pathogens have developed exquisite regulatory systems that not only sense the diverse signals but also trigger specific responses by altering gene expression (1–3). In bacteria, stress adaption is largely achieved by regulation at transcriptional level through the activities of various transcription factors (TFs) (4). Efficient and specific binding of TFs to their targeting promoters is the key for the success of stress adaptation and survival of the pathogens. However, a method to monitor the in situ TF-promoter binding in living bacteria, especially in the complex cellular environment under stresses, is lacking.

Owing to its importance in transcription regulation and stress adaptation, several methods to characterize the TF-promoter interactions, including both in vitro and in vivo, low- and high throughput, have been developed. In vitro methods include the well-established electrophoretic mobility shift assay (EMSA) (5, 6), filter-binding assay (7), DNA foot-printing (8), isothermal titration calorimetry (ITC) (9), and surface plasmon resonance (SPR) (10) etc. that detect the direct binding of TF to its promoters qualitatively and quantitatively. However, a common drawback of these in vitro methods is that the protein and DNA concentrations utilized are usually higher than those in the cell and the DNA fragment is in naked form instead of the compact chromatin state in the cell. Hence, these methods are unable to precisely recapitulate the TF-promoter recognition and binding during the stress adaptation processes in living cells. The high throughput approaches developed in recent years such as chromatin immunoprecipitation coupled with sequencing (ChIP-Seq) and systematic evolution of ligands by exponential enrichment (SELEX) are capable of capturing the TF-promoter binding in vivo occurred under specific culture conditions (11, 12). However, they are generally employed to provide a snapshot of the TF binding sites in the entire genome of bacterial cells and the techniques rely on high throughput sequencing facilities, limiting their widespread applications and accessibilities. Furthermore, all these currently available methods suffer from a limited dynamic range of the TF-promoter interactions detected and are incapable of monitoring the TF-promoter binding elicited by transient or indirect signals present in living bacteria and at the bacteria host interface.

Förster resonance energy transfer (FRET) measurement is an extensively employed strategy to monitor conformational changes and associations between macromolecules in living organisms in real time (13–15). It measures the dipole-dipole energy transfer between two fluorophores (a donor and an acceptor) that are attached to the same biomolecule (intramolecular FRET) or a pair of biomolecules of interests (intermolecular FRET), respectively, when they are in nanoscale proximity and have spectral overlap. Since the efficiency of the energy transfer is reversely correlated to the six power of the distance of the two fluorophores (16), FRET measurement reliably indicates the close proximity and alignment of the macromolecule linker (intramolecular FRET) or macromolecule pairs (intermolecular FRET) onto which the donor and acceptor fluorophores are attached with extreme sensitivity and accuracy. Furthermore, FRET spectroscopy and microscopy represents almost the most simple and accessible form of super-resolved optical measurement (17–19), enabling quantitative and in situ characterization of macromolecule associations both in vitro and in living organisms. Attempts to measure protein-DNA interaction by FRET technique have been reported (20–23). However, majority of these assays were developed for measurement of protein-DNA fragment interactions in vitro due to the simplicity of fluorescent labeling of synthetic DNA fragments and recombinant proteins. Furthermore, intramolecular FRET in which the donor and acceptor fluorophores were dually attached to a single biomolecule was often employed and protein-DNA interaction was inferred by the conformational changes of this fluorescent labeled DNA or protein component (23, 24). These approaches are indirect and inapplicable to monitor the TF-promoter DNA binding in the complex cellular environments in living cells. Intermolecular FRET, on the other hand, is ideal to report the direct interaction of two macromolecules onto which the FRET fluorophore pairs are attached separately. Despite this advantage, an intermolecular FRET-based assay to detect macromolecule interactions in living cells is technically challenging due to the exquisite sensitively of FRET which requires both the close proximity and optimal alignment of the donor and acceptor pairs and the necessity of controlling the ratio of the donor and acceptor fusions to minimize background noise.

In this study, we design an intermolecular FRET-based assay system to monitor TF-promoter binding in living bacteria by employing a pair of bright fluorophores to separately label the TF protein and cellular chromosome DNA. To overcome technical challenges of establishing an intermolecular FRET sensor in live cells, we introduce fluorescence property into TF proteins through genetic encoding of a fluorescent unnatural amino acid (FUAA) at a defined site, i.e., location of TF proteins, i.e., the DNA-binding domain, as the donor fluorophore (25), and employ the cell permeable nucleic acid dye SYTO 9 which is highly selective to DNA molecules (extinction coefficient > 50,000 cm^-1^M^-1^) with a substantial quantum yield (0.58) as the acceptor fluorophore to stain the cellular chromosome DNA. (26). To establish a complementary fluorophore that pairs with SYTO 9, we compared the fluorescence properties of a handful of FUAAs with available aminoacyl-tRNA synthetase (aaRS)-tRNA pairs which are required for genetic incorporation (Table S1) (26–31). An FUAA with the 7-hydroxycoumarin fluorophore, L-(7-hydroxycoumarin-4-yl) ethylglycine (CouA), which is characterized by an emission spectrum peaked at 450 nm with a high quantum yield (0.63) (27) displays the desired properties for spectral overlap with the DNA-bound SYTO 9 (absorption peak ∼500nm). Moreover, the fluorophore affords a large Stokes shift (27) (Fig. S1) which is ideal for intermolecular FRET sensors due to free or minimal cross-excitation. The theoretical Förster radius of this new FRET pair (CouA-SYTO 9) is calculated to be 52.5 Å (Fig. S1) which should allow the occurrence of FRET when the distance between the two fluorophores is in the range of ∼ 25-75 Å.

Herein, we first validated the applicability and efficiency of the new FRET pair CouA-SYTO 9 to report the intricate TF-promoter interactions between a canonical TF protein CueR (CueR-F58CouA) and SYTO 9-stained targeting promoter fragment P*_copA_* in vitro. An intermolecular FRET-based whole-cell assay employing *E. coli* cells expressing CueR-F58CouA in the presence of SYTO 9 stain was then established to measure the CueR-promoter interaction in living bacteria in response to copper (Cu) stress. Specificity of the assay was verified by the abolishment of FRET upon mutating the key arginine residues responsible for promoter binding in CueR or deleting its target promoters P*_copA_* and P*_cueO_* in the chromosome of *E. coli*. Followingly, the assay was extended to monitor the signal transduction and transcription regulation elicited by two-component systems and was employed to identify novel signals that modulate the regulatory systems of interest. Its application to report the TF-promoter binding in *E. coli* colonized in the *C. elegans* host was also demonstrated.

## Materials and Methods

### Construction of plasmid for TF protein expression

DNA fragment encoding the TF protein of interest was obtained by polymerase chain reaction (PCR) amplification using genomic DNA of *E. coli* MG1655 as the template. Primers used are listed in Table S2. The fragment and pET28a plasmid were both digested with NcoI and XhoI for 3 hrs at 37°C. Following purification using the MiniBEST agarose gel DNA extraction kit (TaKaRa), the digested DNA fragment and pET28a were ligated using the quick ligation kit (NEB). The resulting construct was selected on LB agar plates supplemented with kanamycin (20 μg/ml) and was verified by DNA sequencing (BGI, Shenzhen). All plasmids constructed are listed in the supplementary Table S3.

### Amber codon mutation of selected CouA incorporation sites

Each of the selected amino acid sites for CouA incorporation was mutated to a TAG amber codon by site-directed mutagenesis PCR. PCR was conducted using a pair of primers for the desired mutagenesis (Table S2) and pET28a-*cueR*, pET28a-*basR* or pET28a-*phoP* plasmid as the template. PCR product was purified using MiniBEST agarose gel DNA extraction kit (TaKaRa). Following DpnI digestion of the template at room temperature overnight, the PCR product was transformed into *E. coli* DH5α. The transformants were recovered on LB agar plates supplemented with kanamycin (20 μg/ml) at 37°C overnight. Colonies recovered were verified by DNA sequencing (BGI, Shenzhen).

### CRISPR/Cas9-mediated deletion of P_copA_ and P_cueO_ in E. coli chromosome

Deletion of chromosomal *copA* and *cueO* promoter regions were achieved employing a CRISPR/Cas9-mediated approach developed by Zhao et al. (32) with modifications. *E. coli* BL21 (DE3) was electrotransformated with pCAGO plasmid which contains both SpCas9 and λ-RED to obtain BL21-pCAGO. A single colony of the resulting transformant was inoculated into LB medium supplemented with ampicillin (100 mg/ml) and grew overnight at 30°C. 100 μl of the overnight culture was inoculated into 10 ml fresh culture medium and cells were grown at 30°C with IPTG (0.1 mM) induction to OD_600_ as 0.6. The cells were collected and washed with cold ddH_2_O for three times, generating competent cells for electroporation. An aliquot comprising 100 μl competent cells was mixed with 800 ng of the editing-cassette which encompasses 40 bp of upstream-homologous arm of *cueO* promoter region (-74 to -1) followed by the chloramphenicol resistant gene, the N20PAM sequence (GTCCATCGAACCGAAGTAAGG), the other same upstream-homologous arm of *cueO* promoter region, and 40bp downstream-homologous arm, in a 2 mm Gene Pulser cuvette (Bio-Rad) and was subjected to electroporation at 2.3 kV. Following recovering the transformants by cultivation in LB medium for 2 hrs at 30°C, the mixture was spread onto LB agar plate supplemented with ampicillin (100mg/ml), chloramphenicol (25 mg/ml), and 1% glucose (to avoid the background expression of the Cas9 protein). Single colonies were collected and verified by colony PCR to obtain an authentic clone in which the *cueO* promoter region is replaced by the editing-cassette. The confirmed colony was inoculated into LB medium supplemented with ampicillin (100 mg/ml), IPTG (0.1 mM), and L-arabinose (0.2%) at 30°C overnight to enable expression of SpCas9 and the λ-RED recombinase. The cells were then spread on LB agar plates supplemented with ampicillin (100mg/ml). Following colony PCR, desired colonies containing deletion of the promoter of *cueO* gene (AY7037) were verified by DNA sequencing (BGI, Shenzhen). Deletion of the promoter region (-80 to -1) of *copA* gene in AY7038 was conducted following the same procedures except for the editing cassette sequence which encompasses 40 bp of upstream-homologous arm of *copA* promoter region followed by the chloramphenicol resistant gene, the N20PAM sequence, the other same 40 bp upstream-homologous arm of *copA* promoter region, and 40 bp downstream-homologous arm. After obtaining all desired genome modifications, the editing plasmid was cured by growing the resulting cells at 42°C for 48 hrs.

### CouA incorporation in TF proteins and purification of the resulting protein

The pET28a-*cueR*-F58TAG plasmid expressing a C-terminal His_6_-tagged CueR-F58TAG was co-transformed with pEVOL-CouRS that contains tRNA_CUA_ and aminoacyl tRNA_CUA_ synthetase (27) into *E. coli* BL21 (DE3). Transformants were grown on an LB agar plate supplemented with kanamycin (20 μg/ml) and chloramphenicol (25 μg/ml). A single colony of the resulting transformant was inoculated into LB medium supplemented with kanamycin (20 μg/ml) and chloramphenicol (25 μg/ml) and grew overnight at 37°C. 1 ml of the overnight culture was inoculated into 100ml of fresh culture medium and cells were grown at 37°C to OD_600_ as 1.0. Following induction of protein expression with 0.04% arabinose and 0.5 mM IPTG in the presence of 1 mM CouA (Shanghai VastPro Technology) for 12 hrs at 30°C, cells were harvested by centrifugation (4000×g) for 15 min. Cell pellet was resuspended in 10 ml lysis buffer (20 mM Tris-HCl, pH 8.0, 2 M NaCl) and was subject to sonication on ice. Cell debris was removed by centrifugation at 13000×g at 4°C. 1ml Ni-NTA agarose (QIAGEN) was added to the collected supernatant and the mixture was gently shaken at 4°C for 2hrs. Supernatant was removed by gravity column and agarose beads were washed with 50 ml washing buffer (20 mM Tris-HCl, pH 8.0, 2 M NaCl, 30 mM imidazole). CueR-F58CouA protein was eluted using elution buffer (20 mM Tris-HCl, pH 8.0, 2 M NaCl, 500 mM imidazole). Purified protein was subsequently suspended in the buffer for binding assay (20 mM Tris-HCl, pH 8.0, 150 mM NaCl) by buffer exchange through ultrafiltration at 4000×g.

### Verification of CouA incorporation by SDS-PAGE and LC-MS/MS

Purified CueR-F58CouA, BasR-D182CouA, and PhoP-A182CouA protein were analyzed by SDS-PAGE electrophoresis. 15μl of each sample was loaded onto SDS-PAGE (Bio-Rad Mini-PROTEAN Tetra). Following electrophoresis for 60min at room temperature, the gel was imaged under UV transilluminator (UVP) to record fluorescent band and then stained with Coomassie blue. The image of stained gel was recorded by Gel Doc™ XR+ imaging system (Bio-rad). Gel slices corresponding to CueR-F58CouA (16.2 kDa), BasR-D182CouA (26.1 kDa), and PhoP-A182CouA (26.6 kDa) was excised and sent for LC/ESI-MS/MS (Dionex UltiMate 3000 RSLC nano Liquid Chromatography & Orbitrap Fusion Lumos Tribrid Mass Spectrometer).

### TF protein structure prediction (PhoP as an example)

The 3D structure of transcription factor or response regulator protein of two-component systems was predicted via I-TASSER server (https://zhanglab.ccmb.med.umich.edu/I-TASSER/). Amino acid sequences of these proteins were submitted to the server to yield a prediction on the secondary structure, solvent accessibility and normalized B-factor. Ten templates which display the highest significance in the threading alignments from the PDB library were then identified. The server calculated two feasible models, model 1 and model 2 based on the ten templates. The resulting prediction with a higher confidence score (C-score), i.e., model 1 (0.95) in the case of the PhoP protein, was adopted to illustrate the structure of the protein in a PDB file.

### Phosphorylation assay using Phos-tag SDS-PAGE

In vivo phosphorylation of BasR was examined using phos-tag SDS-PAGE following our previous description with minor modifications (33). *E. coli* MG1655 cell harboring pET28a-pBAD-*basR*-FLAG (AY7068) was inoculated into MOPS minimum medium supplemented with kanamycin (20 μg/ml) and grew overnight at 37°C. 50 μl of the overnight culture was inoculated into 5 ml fresh MOPS medium and grew at 37°C for 4 hrs. Expression of BasR was induced with 1mM arabinose for 1 hr. The cells were collected by centrifugation at 4000×g. The cells were washed with MOPS medium supplemented with 0.1, 0.5, or 1 mM FeCl_3_ twice and then cultivated in the corresponding medium for 1hr. 1ml of cells with OD_600_ ∼ 0.5 was collected for the subsequent analysis. The cell was washed with cold PBS twice and resuspended in 200 μl sample buffer (13 mM Tris, 0.2% SDS, 5% glycerol, 0.004% bromophenol blue, pH 6.8) before being subjected to cell lysis by sonication. Following centrifugation at 13000 rpm for 10 min at 4°C, 25 μl of the sample was loaded onto the 10% Tris-Gly polyacrylamide gel containing 25 mM Phos-tag Acrylamide (Wako) and 50 mM MnCl_2_. The samples were separated by electrophoresis for 2 hrs at 4°C. The gel was then washed with 50 ml transfer buffer containing 10 mM EDTA for 20 min to remove residual Mn^2+^. Proteins on the gel were transferred onto a PVDF membrane. The membrane was blocked with 5% non-fat milk in TBST buffer and was incubated subsequently with monoclonal ANTI-FLAG^®^ M2 antibody (Sigma) and goat anti-mouse IgG (H+L)-HRP conjugate (Bio-rad). To detect FLAG-tagged protein bands, the membrane was submerged in the ECL Plus Western Blotting Detection Reagents (GE Healthcare) for 2 min and then subjected to the imaging system (UVITEC Cambridge) to record signals.

### FRET-based CueR-promoter DNA binding assay in vitro

The 185bp promoter region of *copA* gene (P*_copA_*, -178 to +7 bp) was obtained by PCR using genomic DNA of *E. coli* MG1655 as the template. An additional DNA fragment (139 bp) located in the coding region of the *cueR* gene was also obtained by PCR and served as the negative control for the in vitro binding assay. 10 μM purified CueR-F58CouA was mixed with 0.3 μM DNA fragment (P*_copA_* or Frag 1) in 100 μl assay buffer (20 mM Tris-HCl, 150 mM NaCl, pH 8.0) and was incubated at room temperature for 15min. 15uL SYTO 9 (50 μM in ddH_2_O) was then added (final concentration 7.5 μM). Following incubation for 15 min in dark, fluorescence spectrum of the mixture was recorded using Thermo Scientific Varioskan^®^ Flash spectral scanning multimode reader.

### Whole-cell FRET measurement to detect in vivo CueR_CouA_ - promoter_SYTO 9_ binding

A fresh single colony of *E. coli* cell (AY3660) harboring pET28a-*cueR*-F58TAG and pEVOL-CouRS was inoculated into LB medium supplemented with kanamycin (20 μg/ml) and chloramphenicol (25 μg/ml) and grew overnight at 37°C. 75 μl of the overnight culture was inoculated to 7.5 ml fresh medium to subculture the bacteria. Following culturing at 37°C for 4 hrs, CouA incorporation and expression of CueR-F58CouA (or CueR-T27CouA, CueR-Y39CouA, CueR-Q74CouA) was induced mildly by supplementing 0.5 mM IPTG, 0.4% arabinose and 1 mM CouA at 37°C for 4 hrs. Metal ion (stimulating signals) of 0.1 mM CuSO_4_, 0.02 mM MnCl_2_, or 0.02 mM NiCl_2_ (final concentration) was also added. Approximately 8×10^9^ cells were harvested and washed twice with PBS buffer. Following centrifugation at 4000×g for 15 min at 4°C, cell pellet was resuspended in 600 μl of PBS. 85 μl of the suspension was added in each of 6 wells (3 wells for control and 3 wells for experimental group) on a 96-well black plate preloaded with 15 μl of 50 μM SYTO 9 and was incubated for 15 min at room temperature in the dark before being subjected to fluorescence recording.

### In vivo BasR_CouA_ - promoter_SYTO 9_ binding assay

Plasmid pET28a-pBAD-*basR*-D182TAG, G137TAG, E146TAG or S167TAG was co-transformed with pEVOL-CouRS into *E. coli* MG1655 *ΔbasR* strain, resulting in AY7064, AY7065, AY7066 and AY7067 constructs, respectively. Culture conditions for mild protein induction and CouA incorporation were similar to that described for the CueR_CouA_ - promoter_SYTO 9_ binding assay except that 1) the inoculum was cultured in MOPS minimum medium at 37°C for 4hrs before induction; 2) the induction was conducted in the presence of 0.9% arabinose and 1mM CouA at 37°C for 5 hrs. Approximately 3.5×10^10^ cells were harvested and washed twice with fresh MOPS medium. The cells were resuspended in MOPS medium supplemented with 0.5 mM FeCl_3_ as the inducing signal for the BasSR TCS system and shake for 1 hr 37°C. After centrifugation at 4000×g for 15 min at 4°C, cell pellet was resuspended in 600 μl of PBS. The sample preparation and fluorescence spectrum recording were performed as described in the CueR_CouA_ - promoter_SYTO 9_ binding assay.

### In vivo PhoP_CouA_ - promoter_SYTO 9_ binding assay

The pET28a-*phoP*-S137TAG, D145TAG or A182TAG plasmid was co-transformed with pEVOL-CouRS into *E. coli* BL21 (DE3), resulting in AY3707, AY3708, and AY3709 constructs, respectively. Culture conditions for mild protein induction and CouA incorporation were similar to that described for the CueRCouA-promoter DNASYTO 9 binding assay except that 1) the induction was conducted in the presence of 0.5 mM IPTG, 0.4% arabinose and 1 mM CouA at 37°C for 5 hrs, and 2) the stimulants for the PhoPQ system were applied prior to fluorescence signal recording rather than during the induction. Approximately 6×10^9^ cells were harvested and were subjected to sample preparation. Following incubating 85 μl of cell suspension containing mildly induced PhoP-CouA protein and 15 μl of SYTO 9 (50 μM) at room temperature for 15 min in the dark, different stimuli was added in the buffer. 1mM of

MgCl_2_ was added to the cell suspension before adding other stimulants. EDTA, polymyxin B, tryptophan, indole, IAA, gentamycin was added at the designated concentrations in the text. The suspension is subjected to fluorescence spectrum recording as described above. Zero-time is defined as the fluorescence of the assay system recorded immediately following the addition of stimulants.

### Fluorescence spectroscopy and FRET effect quantification

Fluorescence spectra of the assay suspension described above were recorded using Thermo Scientific Varioskan^®^ Flash spectral scanning multimode reader at 25°C. The excitation wavelength was fixed at 360 nm, and the scanning wavelength was recorded from 400 to 550 nm. Degree of the FRET effect was quantified by the ratio of

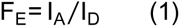

where *I*_*A*_ is the fluorescence intensity of CouA at 500 nm, and *I*_*D*_ is the fluorescence intensity at 450 nm.

The Förster radius *R*_0_ is the Förster distance which is the distance between the FRET donor and acceptor when the energy transfer efficiency is 50%. *R*_0_ is calculated from the equation (34):

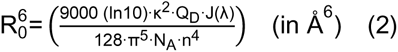

Where *k*^2^ is the orientation factor in resonance energy transfer, *Q*_*D*_ is the quantum yield of the donor in the absence of acceptor, *N*_*A*_ is Avogadro’s number, *n* is the refractive index of the medium, and *j*(*ƛ*) is the overlap integral. To calculate this equation, *k*^2^ is designated as 2/3, *Q*_*D*_ as 0.63, n as 1.4, and *N*_*A*_ as 6.02214076×10^23^. The *J*(*ƛ*) is calculated by (34):

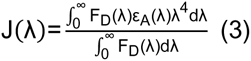

where *F*_*D*_ is the emission spectrum of the donor, *ε*_*A*_(*ƛ*) is the molar extinction coefficient of the acceptor at *ƛ*, which is 50000 cm^-1^M^-1^ for SYTO 9.

### Preparation of E. coli cells expressing PhoP-A182CouA and C. elegans *DA597* for fluorescent microscopic measurement of FRET

*C. elegans phm-2*(-) mutant strain DA597, which lost the grinder function, was obtained from the Caenorhabditis Genetics Center (CGC), which is supported by the National Institutes of Health - Office of Research Infrastructure Programs (P40 OD010440). *E. coli* BL21 cells expressing PhoP-A182CouA (AY3706) was cultured in LB medium supplemented with kanamycin (20 μg/ml) and chloramphenicol (25 μg/ml) at 37°C for 5 hrs in the presence of 0.5 mM IPTG, 0.4% arabinose and 1 mM CouA. Cells were collected and washed with M9 buffer (0.3% KH_2_PO_4_, 0.6% Na_2_HPO_4_, 0.5% NaCl, 1 mM MgSO_4_). 5 μl of 50 μM SYTO 9 was added in 1 ml cell suspension (final concentration 0.25 μM), which was then incubated for 15 min in the dark to enable the entry of SYTO 9 into *E. coli* cells. The suspension was washed with 1 ml M9 buffer for three times and finally resuspended in 100 μl of M9 buffer and was then placed in the center of a 60 mm NGM agar plate (0.25% peptone, 0.3% NaCl, 1.7% Bacto-agar, 1 mM CaCl_2_, 1 mM MgSO_4_, 5 mg cholesterol, 25 mM phosphate buffer) in a circle (20 mm diameter) as food for the *C. elegans* DA597 strain. This plate is termed as AY3706-NGM plate. The eggs of DA597 were then seeded onto NGM agar plate containing *E. coli* OP50 (OP50-NGM) and fed for 48 hrs at 20°C. Ten L4 stage larva grown on the OP50-NGM plate were then transferred to the AY3706-NGM plate and fed with the AY3706 bacteria for 2 hrs at 20°C before being subjected to fluorescent microscopy imaging.

### FRET recording using confocal fluorescent microscope

The larva grown on the AY3706-NGM plate described above following 2 hr-incubation were placed onto an agarose pad containing 3% agarose and 4% 2,3-butanedione monoxime on a microscope slide. Fluorescence signal emitted from the larva was recorded using Zeiss LSM 710 confocal microscope using 40× oil immersion objective lens. The sample was excited at 405 nm and the emission fluorescence of CouA was recorded from 420 nm to 480 nm. To record the emission fluorescence of SYTO 9, the sample was excited at 488 nm and emission fluorescence was recorded from 490 nm to 600 nm. To record the fluorescence effected by FRET, the sample was excited at 405 nm by laser and emission fluorescence was recorded from 490 nm to 600 nm.

## Results

### CouA is successfully incorporated into a canonical TF protein CueR at a defined site

To establish an intermolecular FRET-based transcription factor (TF)-promoter binding assay based on intermolecular FRET, we first explored the site-specific incorporation of CouA into the DNA binding domain of a canonical TF protein CueR which displays intricate interactions with its targeting promoters during the switch from a transcription repressor to an activator in response to the inducing signal. CueR is a copper (Cu) and silver (Ag) metalloregulatory MerR family protein that senses Cu(I) stress and activates the expression of two target genes, *copA* and *cueO*, which encodes the ATPase copper exporter CopA and copper oxidase CueO, respectively (36–38) (Fig. 1A). The mechanism of transcription regulation by CueR has been elucidated by both crystallography in which CueR is bound with a 23-bp promoter DNA (39) and by cryo-EM in which CueR, RNAP and a 58-bp promoter DNA forms a complex (40). These structural studies have suggested that CueR binds to target promoters and activates gene transcription by allosteric DNA distortion. Since CouA contains an aromatic sidechain, we first selected aromatic amino acid residues located at the relatively unstructured loop regions in the DNA binding domain (DBD) of CueR as the potential sites for CouA incorporation. The DBD of CueR is composed of four α-helixes and is connected to a two-turn C-terminal α-helix by a hinge loop and a long dimerization helix (39). It was shown that CueR binds to its target promoter as a repressor in the absence of the inducing signal Cu (39). Upon being bound by Cu(I), CueR undergoes a conformational change and is switched to a transcription activator, resulting in a twist of the bound DNA by tightly clamping the α2 and a loop wing of the DBDs, i.e., R18, 31, and 37, into DNA major grooves (Fig. 1A) (39). Based on this structural information, we chose F58 which is located at the loop region between the helix α3 and helix α4 for CouA incorporation (Fig. 1A). The calculated distance between the selected site to the two closest DNA minor grooves which should be bound by the SYTO 9 dye is ∼22 Å and 45 Å, respectively. These distances are within the calculated distance range (25-75 Å) for FRET occurrence based on the theoretical Förster radius of the CouA-SYTO 9 pair (52.5 Å) (Fig. S1).

**Figure 1.**
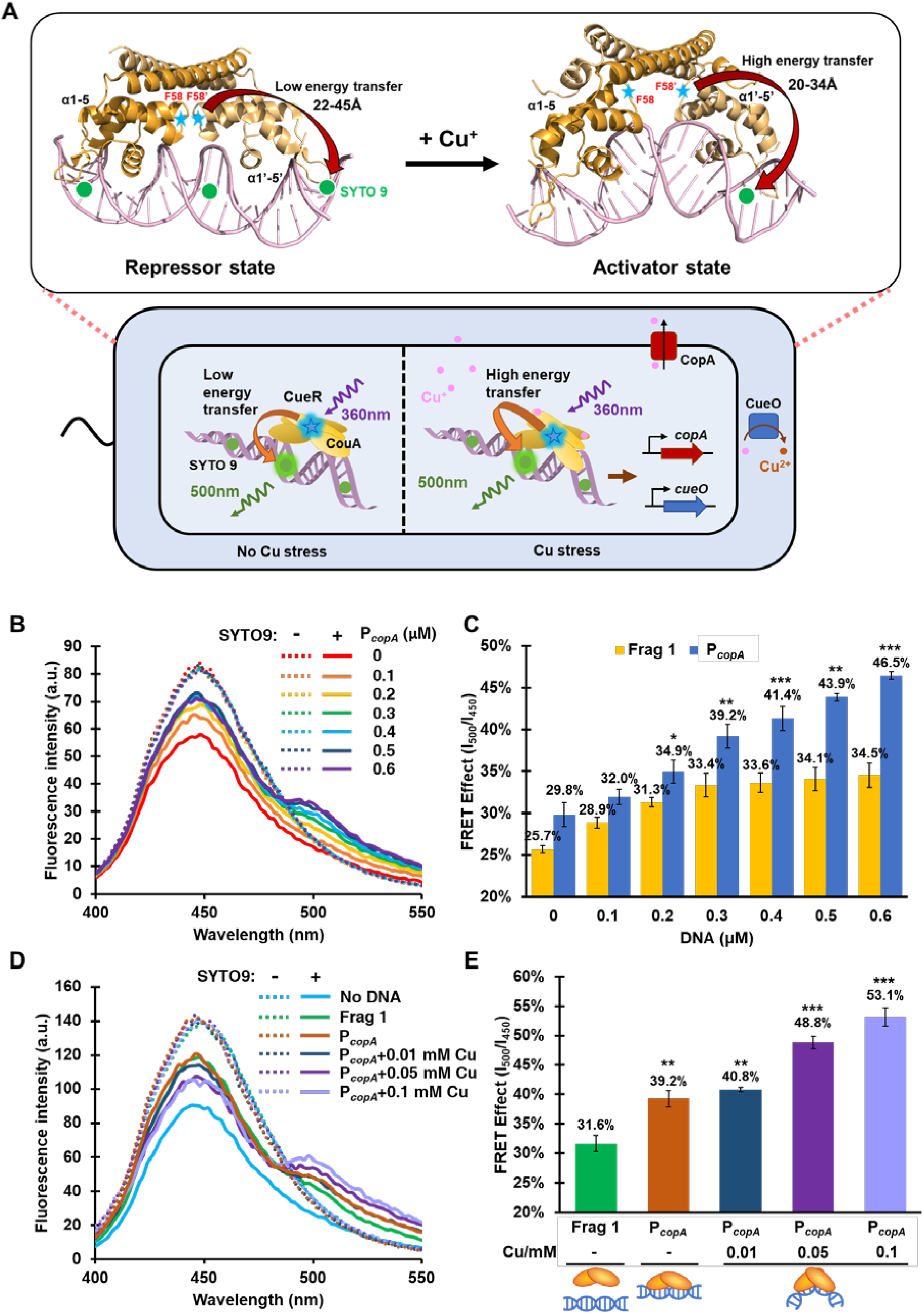
Schematic diagram of the intermolecular FRET-based TF-promoter binding assay and verification of the CouA-SYTO 9 FRET pair employing the activities of CueR as an example. (A) Diagram of the intermolecular FRET-based TF-promoter binding assay. Structure of the CueR-promoter DNA complex in the repressor and activator state is shown. Ribbon structure of CueR-DNA complexes were adopted from the crystal structure of *E. coli* CueR bound to the *copA* promoter DNA (PDB ID: 4WLW, 4WLS). The site for CouA incorporation, F58, was shown. Calculated distance between CouA in the F58 position and SYTO 9 bound to a proximal DNA minor groove was shown. FRET effect in the activator-promoter binding state (increase in FRET efficiency) is expected. Blue spheres, CouA (Ex: 360 nm, Em: 450 nm); Green spheres, SYTO 9 (Ex: 483 nm, Em: 503 nm). (B) Fluorescent spectra of 10 μM CueR-F58CouA purified protein mixed with the P*copA* DNA fragment (0-0.6 μM) in the presence of SYTO 9 (7.5 μM). (C) FRET effect (reflected by intensity ratio of 500 nm to 450 nm) of the CouA-SYTO 9 pair. (D) Fluorescence spectra of purified CueR-F58CouA incubated with a random DNA fragment (Frag 1), P*copA*, or P*copA* with a series concentration of Cu^+^ (0-0.1 mM) in the presence or absence of SYTO 9. (E) FRET effect of the CouA-SYTO 9 pair in the assay mixtures. P*copA*, DNA fragment of promoter region of *copA* gene; Frag1, random DNA fragment corresponding to +1 - +142 bp in CueR. Data are the mean of three biological repeats and are expressed as mean ± SD. *, *P*<0.05; **, *P*<0.01; ***, *P*<0.001 (based on Student’s *t* test). See also Figure S1, S2 & S3.

The genetic code encoding F58 in CueR was mutated to the amber codon TAG. In the presence of the previously optimized orthogonal tRNA/tRNA acetyltransferase MjtRNA-MjTyrRS (27) and the supplement of 1mM CouA (Fig. S2A), a protein band (16.2 kDa) corresponding to CueR-F58CouA-His_6_ was detected by fluorescent SDS-PAGE (Fig. S2B). The protein was subsequently purified and was subjected to LC-MS/MS analysis. Site-specific incorporation of CouA in CueR by replacing F58 was verified by the expected mass of y26 and b4-b15 peptides with a 174.15 Da addition relative to the corresponding peptides derived from the WT CueR (Fig. S2C).

### FRET effect between CueR-F58CouA and SYTO 9-labeled P_copA_ reliably reports the CueR-P_copA_ binding

We next examined the occurrence of FRET and its degree effected by the binding of purified CueR-

F58CouA with the 200-bp DNA fragment of P*_copA_* stained with SYTO 9 *in vitro*. Emission spectra of the

CueR-F58CouA protein, CueR-F58CouA with the P*_copA_* DNA fragment, and CueR-F58CouA with the free SYTO 9 were firstly recorded. Comparing with the spectra of CueR-F58CouA alone (red dash line, Fig. 1B) and CueR-F58CouA mixed with varying amounts of P*_copA_* (dash lines, Fig. 1B) which exhibit the characteristic bright fluorescence of CouA peaked at 450 nm, addition of free SYTO 9 caused a slight decrease of the overall fluorescence intensity (red solid line, Fig. 1B), which was found to be saturated when SYTO 9 concentration is above 5 μM (Fig. S3A). Since the decrease of fluorescence intensity at 450 nm in the mixture was not accompanied by an increase of fluorescence intensity at 500 nm, which is the desired FRET signal for the CouA-SYTO 9 pair, this background change is mostly due to a fluorescence quenching of CouA by the free SYTO 9 and is not a false positive FRET. When the protein (10 μM) was mixed with varying amount of P*_copA_* DNA fragment in the presence of SYTO 9 (7.5 μM), a significant fluorescence emission peaked at 500 nm appeared and the intensity increased in a P*_copA_* concentration dependent manner (Fig. 1B & C), indicating that a FRET effect occurred between CueR-F58CouA and the SYTO 9 bound-P*_copA_*, which led to the energy transfer from CouA to SYTO 9. In contrast, supplying a random DNA fragment (Frag 1) only caused a very moderate increase at 500 nm and the intensity was not increased continuously with the increase of the DNA concentration (Fig. 1C), suggesting that the FRET signal observed in the mixture of CueR-F58CouA and P*_copA_* was effected by the specific interaction between CouA-F58CouA and the SYTO 9 bound-P*_copA_*. This result is consistent with the fact that CueR binds to its target promoters as a repressor in the absence of the inducing signal Cu(I). Since the background-level FRET between CueR-F58CouA with the random DNA was saturated when its concentration was above 0.3 μM, 0.3 μM DNA fragment is applied in the following assays. Next, we supplied the stimulant Cu(I) which is generated by CuSO_4_ in the presence of ascorbate to the assay mixture. A dose-dependent increase of the FRET signal was observed (Fig. 1D & E), indicating a closer proximity and improved alignment between the FRET pairs upon the binding of Cu(I) to CueR which is consistent with the clamp of the DBD into DNA grooves by the Cu(I) bound CueR activator revealed by the structural studies (39). These results demonstrated that the FRET occurrence and its degree between CouA and SYTO 9 reliably reports the intricate interaction of CueR and its targeting promoter DNA in response to the presence and strength of the inducing signal of the regulatory system.

### Whole-cell FRET between CueR_CouA_ and promoter_SYTO 9_ monitors CueR-promoter interaction in vivo

We next employed the FRET pair to monitor the TF-promoter interactions in living *E. coli*. To achieve this, we first optimized the condition for CouA incorporation in *E. coli* cells such that CouA is incorporated into CueR but the protein is not over-expressed to artificially shift or saturate the TF-promoter binding. We carried out CouA incorporation and protein expression under a mild condition (0.5 mM IPTG, 0.4% arabinose, 1 mM CouA for 4 hrs at 37°C) compared to that for over-expression and purification of CueR-F58CouA. The expression level of CueR-F58CouA protein in the cell cultured under this condition was estimated to be ∼113 μg/ml by comparing with a series of purified CueR-F58CouA protein with known concentrations (Fig. S4A) which is less than 1/2000 of total proteins in *E. coli* grown in LB medium at stationary stage (200-320 mg/ml) (41). This expression level affords a fluorescence intensity reading in the range of 80-120 a.u. when 10 O.D. cell suspension was subjected to fluorescence recording. Fluorescent SDS-PAGE (Fig. S2B) and fluorescent microscopy (Fig. S5A) confirmed the expression of CueR-F58CouA and the bright fluorescence of the resulting cell suspension. To determine the concentration of SYTO 9 to be applied in the whole-cell FRET assay, we first examined the background fluorescence intensity of the *E. coli* BL21 cell suspension host stained with a series concentrations of SYTO 9. A concentration dependent increase of the fluorescence intensity at 500 nm was observed in the range of 0-20 a.u. (Fig. S3B). The same SYTO 9 concentration (7.5 μM) as applied in the in vitro assay was selected which is also consistent with the concentration suggested by McGoverin et al. (42) for general application of SYTO 9 to several common bacteria. Staining of the cell suspension with this concentration of SYTO 9 yielded the desired bright green fluorescence (Fig. S5A). These results confirmed the applicability of the two fluorophores in live cells.

A whole-cell FRET measurement to monitor in vivo TF-promoter binding was subsequently established. 0.1 mM CuSO_4_ was added in the cell culture during CouA incorporation and CueR-F58CouA expression to activate CueR-mediated transcription regulation. Following incubation with SYTO 9 for 15 min in the dark, the whole-cell suspension was directly subjected to emission fluorescence spectrum recording (Fig. 2A). Comparing with the spectrum without the Cu stress (cyan line, Fig. 2B) which exhibited a background level FRET (50.8%) (Fig. 2F), an obvious FRET enhancement with a quantified FRET effect of 61.7% was observed in the cell treated with 0.1 mM Cu (purple line, Fig. 2B), suggesting that the CouA-SYTO 9 FRET pair is applicable to report the binding of CueR with its promoter in live *E. coli* cells in response to the Cu stress. To confirm that the observed FRET effect and its enhancement was elicited by the specific binding of CueR with its target promoters (P*_copA_* and P*_cueO_*) in *E. coli* cells, we mutated the key arginine residues R18 and R37 (37) located at the loop wing of the promoter binding domain of CueR which are essential for DNA clamping, and conducted the assays employing *E. coli* cells expressing CueR-F58CouA-R18A and CueR-F58CouA-R37A, respectively. No FRET enhancement was observed when cells expressing either of the mutant proteins were treated by Cu compared to that without Cu treatment (Fig. 2C-D). Quantitation of the degree of FRET confirmed the insignificant FRET changes for CueR-F58CouA-R18A and CueR-F58CouA-R37A as opposed to that (10.9%) for the CueR-F58CouA (Fig. 2F). To further confirm the specificity of the FRET signals observed, we also conducted the assay in a mutant strain in which the two targeting promoters of CueR, P*_copA_* and P*_cueO_*, were deleted from the *E. coli* BL21 genome, i.e., BL21 (DE3)-*ΔP_copA_ΔP_cueO_*. Whole-cell FRET assay in this mutant showed that deletion of the targeting promoters completely abolished the FRET effect between CueR_F58CouA_ and promoter DNA_SYTO 9_ (Fig. 2E & F). These results demonstrated the specificity and sensitivity of the intermolecular FRET-based whole-cell TF-promoter binding assay we developed. To establish the CouA incorporation locations applicable for the intermolecular FRET-based TF-promoter binding assay, we incorporated CouA into several additional locations (Fig. 3A) and constructed the following variants: CueR-T27CouA, CueR-Y39CouA, and CueR-Q74CouA, in which CouA was incorporated into the loop region between the α2 and α3 in the DBD (T27CouA and Y39CouA) and the loop region between the α4 in the DBD and the α5 dimerization helix (Q74CouA). The calculated distances between these sites and the closest central minor grooves which should be bound by SYTO 9 are 26 Å, 30 Å, and 36 Å, respectively. Whole-cell assay showed that an enhancement of FRET upon Cu treatment was also observed in cells expressing CueR-T27CouA (Fig. 3B & E) and CueR-Y39CouA (Fig. 3C & E), but not in cells expressing CueR-Q74CouA (Fig. 3D & E), suggesting that the donor fluorophore (CouA) can be incorporated into multiple locations in the DBD, especially the loop regions between α helices, but not the distal dimerization domain for the FRET-based assay. Notably, we also constructed a plasmid encoding the CueR-P73CouA variant, but the protein was not successfully expressed despite its close proximity to Q74. This result suggested that certain residues, such as proline, is not tolerant to replacement by CouA.

**Figure 2.**
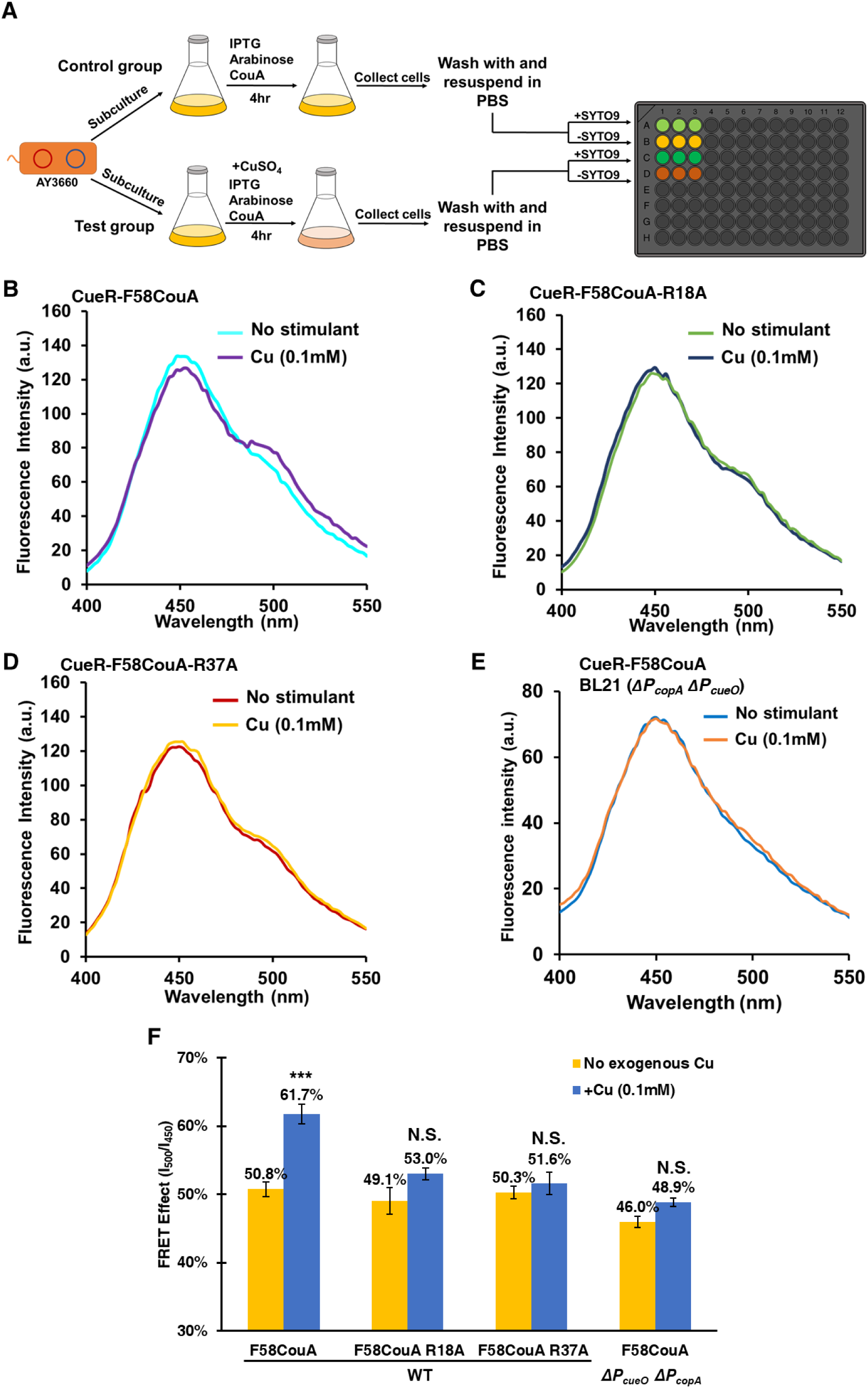
The intermolecular FRET-based TF-promoter binding assay detects the specific interaction of CueR with its regulated promoters in living *E. coli*. (A) Sample preparation workflow for the FRET-based CueR-promoter binding assay in vivo. 0.1 mM CuSO4 was added to the test group to activate CueR. Cell suspension was incubated in the dark for 15 min before being subjected to fluorescence recording. (B-D) Fluorescence spectra of BL21 (DE3) cells expressing CueR-F58CouA (AY3660) (B), or CueR-F58CouA-R18A (AY3719) (C), or CueR-F58CouA-R18A (AY3720) (D), under non-inducing condition and in the presence of 0.1 mM CuSO4. All data are normalized against the conditions of no SYTO 9 addition. (E) Fluorescence spectrum of BL21 (DE3)-*ΔPcopA ΔPcueO* cells (AY7038) expressing CueR-F58CouA under non-inducing condition (blue) and in the presence of 0.1 mM CuSO4 (orange). (F) FRET effect of the CouA-SYTO 9 pair in the designated assay mixtures. Data are the mean of three biological repeats and are expressed as mean ± SD. *, *P*<0.05; **, *P*<0.01; ***, *P*<0.001, N.S., not significant relative to the no metal treatment (based on Student’s *t* test). See also Figure S3, S4 & S5.

**Figure 3.**
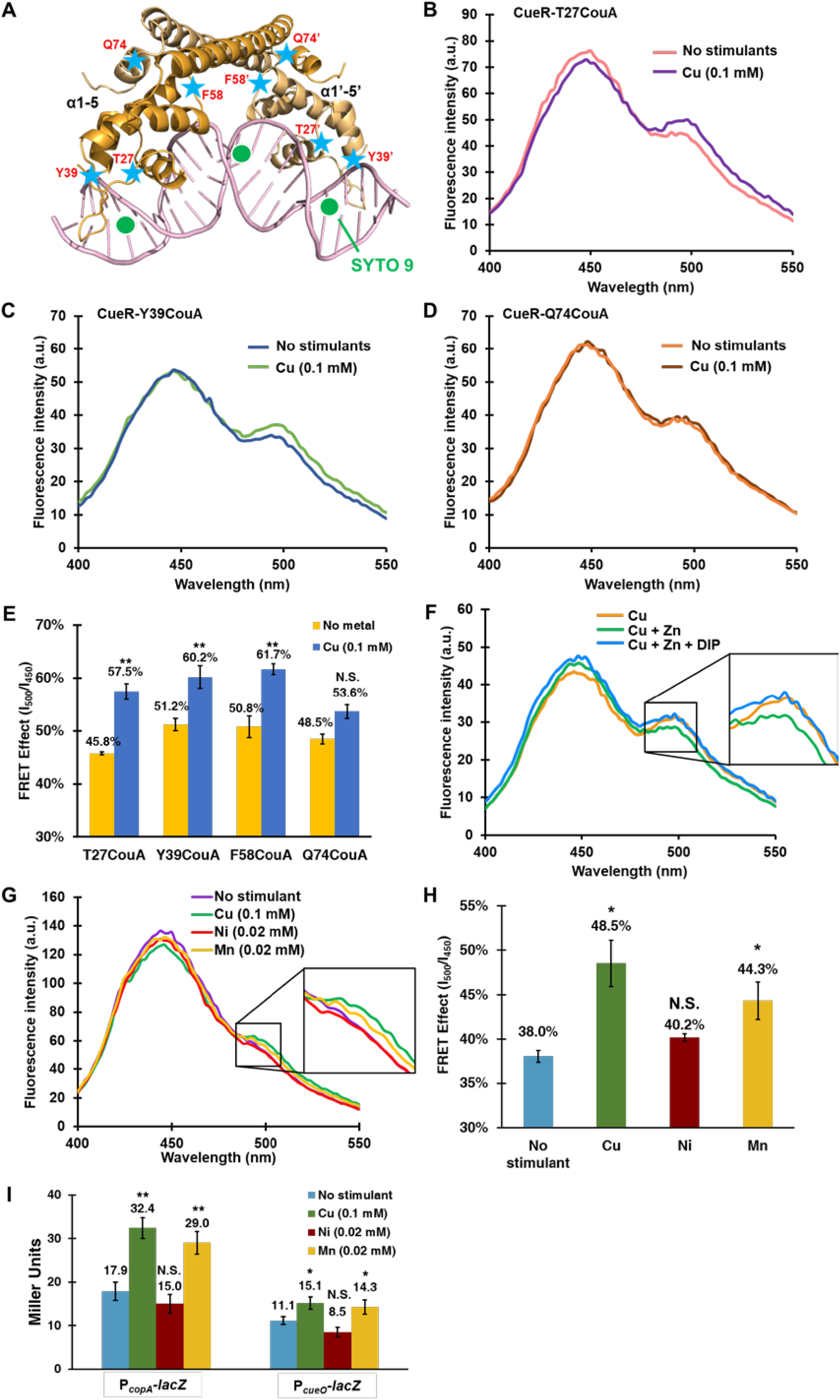
Whole-cell intermolecular FRET-based TF-promoter binding assay reports the intricate interactions of CueR-promoters in living cells. (A) Diagram showing the locations of CouA incorporation sites tested. CueR is shown in its active state (4WLW). The blue star represents CouA, and the green spheres represent SYTO 9. (B-D) Fluorescence spectra of BL21 (DE3) cells expressing CueR-T27CouA (B), or CueR-Y39CouA (C), or CueR-Q74CouA (D) in the presence or absence of 0.1 mM CuSO4. (E) FRET effict of the CouA-SYTO 9 pair in the designated cell suspensions. (F-G) Fluorescence spectra of BL21 (DE3) cells expressing CueR-F58CouA in the presence of 0.1 mM CuSO4 (orange), 0.1 mM CuSO4 and 0.1 mM ZnCl2, and 0.1 mM CuSO4, 0.1 mM ZnCl2 and 0.2 mM DIP (blue) (F), 0.1 mM CuSO4, 0.02 mM NiCl2 (red), and 0.02 mM MnCl2 (dark green) (G). All data are normalized against the conditions without SYTO 9 addition. (H) FRET effect of the CouA-SYTO 9 pair in the designated cell suspensions. (I) β-galactosidase activities of P*copA*-*lacZ* and P*cueO*-*lacZ* transcription fusions in the presence of Cu, Mn and Ni. Data are the mean of three biological repeats and are expressed as mean ± SD. *, *P*<0.05; **, *P*<0.01; ***, *P*<0.001, N.S., not significant relative to no metal treatment (based on Student’s *t* test). See also Figure S8.

### The CouA-SYTO 9 FRET pair reports the intricate CueR-promoter interaction elicited by transient signals in living cells and novel signals that activate CueR

Next, we applied the assay to detect the inactivation of the CueR regulatory system and the activation of the system by indirect or intermediate stimulant. Zinc (Zn) was indicated to inactivate CueR indirectly through transiently elevating intracellular iron (Fe) (43). To verify this regulatory circuit, we examined FRET occurrence and its degree in *E. coli* cells expressing CueR-F58CouA and stained with SYTO 9 in the presence of Zn which should suppress the CueR-promoter interaction, and Zn in combination with the iron chelator 2,2’-dipyridyl (DIP) which should restore the CueR-promoter interaction. As shown in Figure 3F, while a significant FRET was observed in the presence of Cu, addition of Zn abolished the FRET effect. Further supplementing DIP restored the FRET effect, confirming the regulatory circuit of Zn-Fe-CueR (43).

We also employed the FRET-based assay to explore novel signals that can activate CueR. We found that supplementing 0.02mM manganese also resulted in an enhancement of FRET (Fig. 3G) with a calculated FRET change of 6.3% (Fig. 3H), whereas Ni (0.02 mM) did not elicit a FRET enhancement in cells expressing CueR-F58CouA, suggesting that Mn may also serve as a signal to activate CueR. To verify this finding, we performed β-galactosidase activities assay to examine the transcription of *P_copA_-lacZ* and *P_cueO_-lacZ* in the presence of the same concentration of these metals which do not inhibit the growth of *E. coli* cells. β-galactosidase activities assay showed that Mn supplement indeed activated transcription of the *P_copA_-lacZ* and *P_cueO_-lacZ* fusions that are controlled by CueR, whereas Ni supplement did not (Fig. 3I). This result verified that the whole-cell intermolecular FRET-based assay we established is applicable to report TF-promoter interaction occurred in a complex regulatory circuit induced by indirect stimulants and can be employed to identify novel signals and conditions that modulate the regulatory system of interest.

### The intermolecular FRET-based TF-promoter binding assay monitors the signal transduction of two-component regulatory systems

We next extended the methodology to another important type of gene regulatory system in bacteria: the two-component system (TCS). TCS is composed of a membrane-bound histidine kinase (HK) that senses the extracellular signals and a cognate response regulator (RR) that regulates gene transcription in response to signals perceived by the HK. We chose the BasSR TCS to test the application of the whole-cell intermolecular FRET-based TF-promoter binding assay. In this TCS, BasS is the HK which senses high iron (Fe^3+^) level and phosphorylates the cognate RR BasR (Fig. 4A) (44). Phosphorylated BasR binds to the promoters of its target genes *ugd*, *eptA,* and *arn* operon which encodes enzymes that modify lipid A on the outer membrane of Gram-negative bacteria and activates their transcription (Fig. 4A) (45).

**Figure 4.**
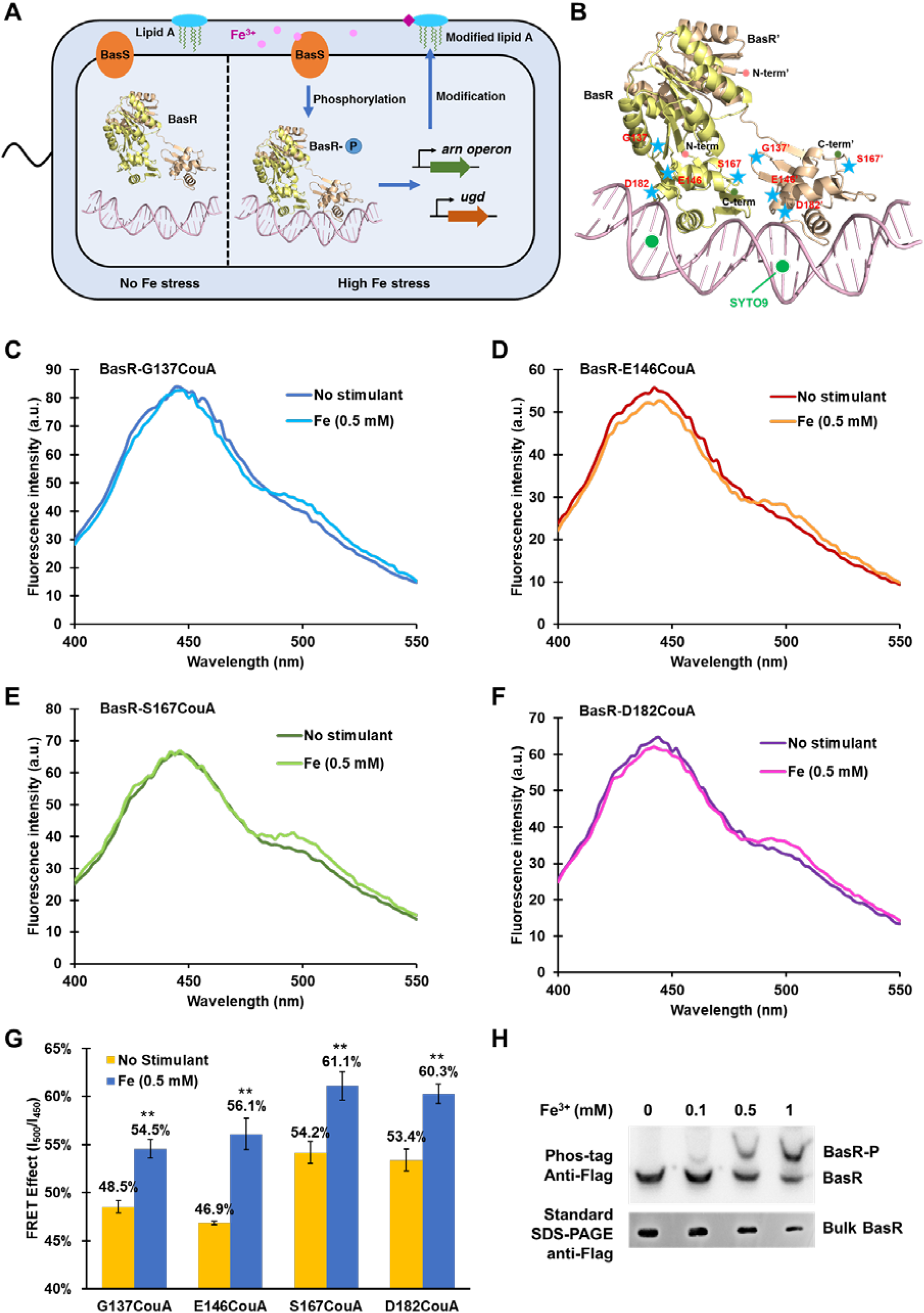
The intermolecular FRET-based TF-promoter binding assay system monitors activities of the two-component system BasSR in vivo. (A) Diagram showing the signal transduction of the BasSR TCS. (B) Diagram showing the locations of CouA incorporation sites in BasR being teste. BasR is shown in its active state. The blue stars represent CouA, and green spheres represent SYTO 9. Ribbon structure of BasR was simulated using the I-TASSER server. DNA fragment was adopted from Crystal structure of *Klebsiella pneumoniae* PmrA (PDB ID: 4S05). (C-F) Fluorescence spectra of MG1655 cells expressing BasR-G137CouA (C), BasR-E146CouA (D), BasR-S167CouA (E) and BasR-D182CouA (F) in the presence or absence of 0.5 mM Fe^3+^. (G) FRET effect of the CouA-SYTO 9 pair in the designated assay mixtures. (H) Phosphorylation of BasR in the presence of a series of FeCl3 detected by Phos-tag SDS-PAGE. Bands corresponding to phosphorylated (BasR-P) and unphosphorylated BasR (BasR) are indicated. Data are the mean of three biological repeats and are expressed as mean ± SD. *, *P*<0.05; **, *P*<0.01; ***, *P*<0.001, N.S., not significant relative to no metal treatment (based on Student’s *t* test). See also Figure S2, S4, S5 & S8.

To monitor the activation of BasR in response to Fe^3+^ stress and interaction of BasR with its target promoters in vivo, we set out to incorporate CouA into the DNA binding domain of the BasR protein. The crystal structure of BasR is not available. To determine the CouA incorporation sites, we employed I-TASSER to simulate the 3D structure of BasR using the crystal structure of its homologue, *Klebsiella pneumoniae* PmrA (PDB ID: 4S05), as the template (46–48). Two domains were identified in the predicted structure, the receiver domain (1-121 AA) and the DNA-binding domain (125-219 AA). The DBD encompasses three α-helices (α6-α8) and two β-sheets (β2, β3). Four potential CouA incorporation sites were selected, G137 (located in β2), E146 (located in a loop region between β2 and α6), S167 (located in a loop region between α6 and α7), and D182 (located in a loop region between α7 and α8) (Fig. 4B). They display a calculated distance of 26 Å, 28 Å, 24 Å, 22 Å, respectively, to the closest DNA minor grooves. Herein, we replaced the T7 promoter in pET28a to pBAD and conducted the assay in the *E. coli* reference strain MG1655 (35). Expression of BasR and CouA-incorporation was carried out in the presence of 0.9% arabinose. Expression of the resulting CouA-incorporated proteins (BasR-S137CouA as an example) was monitored by western-blot (Fig. S4B) and verified by fluorescent SDS-PAGE (Fig. S2B), fluorescent microscopy (Fig. S5B), and LC-MS/MS (Fig. S2D).

To monitor the activation of the BasSR in response to Fe^3+^ stress, *E. coli* MG1655 cells expressing CouA-incorporated BasR was treated with 0.5 mM Fe^3+^ in the MOPS medium for 1hr before being stained with SYTO 9 and subjected to emission fluorescence recording. The whole-cell suspension contained a similar ratio of the donor fusion and acceptor stain as employed in the case of CueR-promoter assay, i.e., 10 O.D cells stained with 7.5 μM SYTO 9. Emission fluorescence recording showed that upon Fe^3+^ treatment, a statistically significant FRET enhancement relative to the non-treatment control was observed in all these cells, i.e., BasR-G137CouA, BasR-E146CouA, BasR-S167CouA, BasR-D182CouA (Fig. 4C-F). The degree of FRET enhancement was calculated to be 6.0%, 9.2%, 6.9% and 6.9%, respectively (Fig. 4G). To verify that the FRET assay indeed reported the activation of BasR and its binding to the target promoters, we examined the phosphorylation of BasR protein under this culture and assay condition using Phos-tag SDS-PAGE. It was shown that a significant portion of BasR was phosphorylated in the presence of 0.5 mM Fe^3+^, confirming that the intermolecular FRET-based assay we developed is applicable to monitor the activation of the TCS in vivo. Furthermore, the fact that incorporation of CouA into all four selected sites resulted in the desired FRET enhancement upon stimulation and that these CouA incorporation sites were determined based on the simulated 3D structure of BasR suggested that the assay system we developed is robust to monitor the diverse signal transduction systems.

### Whole-cell FRET measurement monitors the activation of PhoPQ TCS in response to different stimulants

We next expanded the assay to the PhoPQ regulatory system which is one of the most important TCS in enterobacteria that regulates the virulence and antibiotic resistance in response to the environment of their mammalian hosts, such as low divalent cations, low pH, antimicrobial peptides and osmotic upshift (49–55). In this TCS, PhoQ is the histidine kinase that phosphorylates and activates the response regulator PhoP upon perceiving stimulating signals. Phosphorylated and activated PhoP then binds to the target promoters to regulate gene transcription (Fig. 5A). Since the crystal structure of *E. coli* PhoP is not available, we simulated its 3D structure using the crystal structure of *Mycobacterium tuberculosis* PhoP-DNA complex (PDB ID: 5ED4) as the template. Two domains were shown in the simulated structure, the receiver domain (1-123 AA) and the DBD (123-223 AA). The DBD encompasses three α helixes (α6-α8) and two β-sheets (β2, β3). We selected three potential sites in the DBD for CouA incorporation, S137 (located in β2), D145 (located at the loop region between β2 and α6), and A182 (located at the loop region between α7 and α8) (Fig. 5B). The calculated distances of D145, S137 and A182 to the closest DNA minor groove is approximately 30 Å, 25 Å, and 20 Å, respectively. Plasmids encoding the three proteins PhoP-S137CouA, PhoP-D145CouA, and PhoP-A182CouA were successfully constructed. Fluorescent SDS-PAGE (Fig. S2B), fluorescent microscopy (Fig. S5C), and LC-MS/MS (Fig. S2E) analysis confirmed the expression of these proteins (PhoP-A182CouA as an example) and their fluorescence in *E. coli* cells. We first conducted the whole-cell FRET assay to monitor the activation of the regulatory system under the condition of Mg^2+^ depletion. 10mM Mg^2+^ was added in the cell suspension to ensure that PhoPQ is fully inactivated prior to applying the stimulants as reported by Bader et al. (Bader et al., 2005). Depletion of Mg^2+^ was initiated subsequently by supplying 20 mM EDTA following a previous description (55). An enhancement of FRET was observed in all three assay mixtures relative to the no EDTA control (Fig. 5C-E). Among them, PhoP-A182CouA displayed the greatest degree of FRET effect (7.7%) (Fig. 5F), consistent with the closest calculated distance between this site with the DNA minor groove among the three selected sites in the simulated structure.

**Figure 5.**
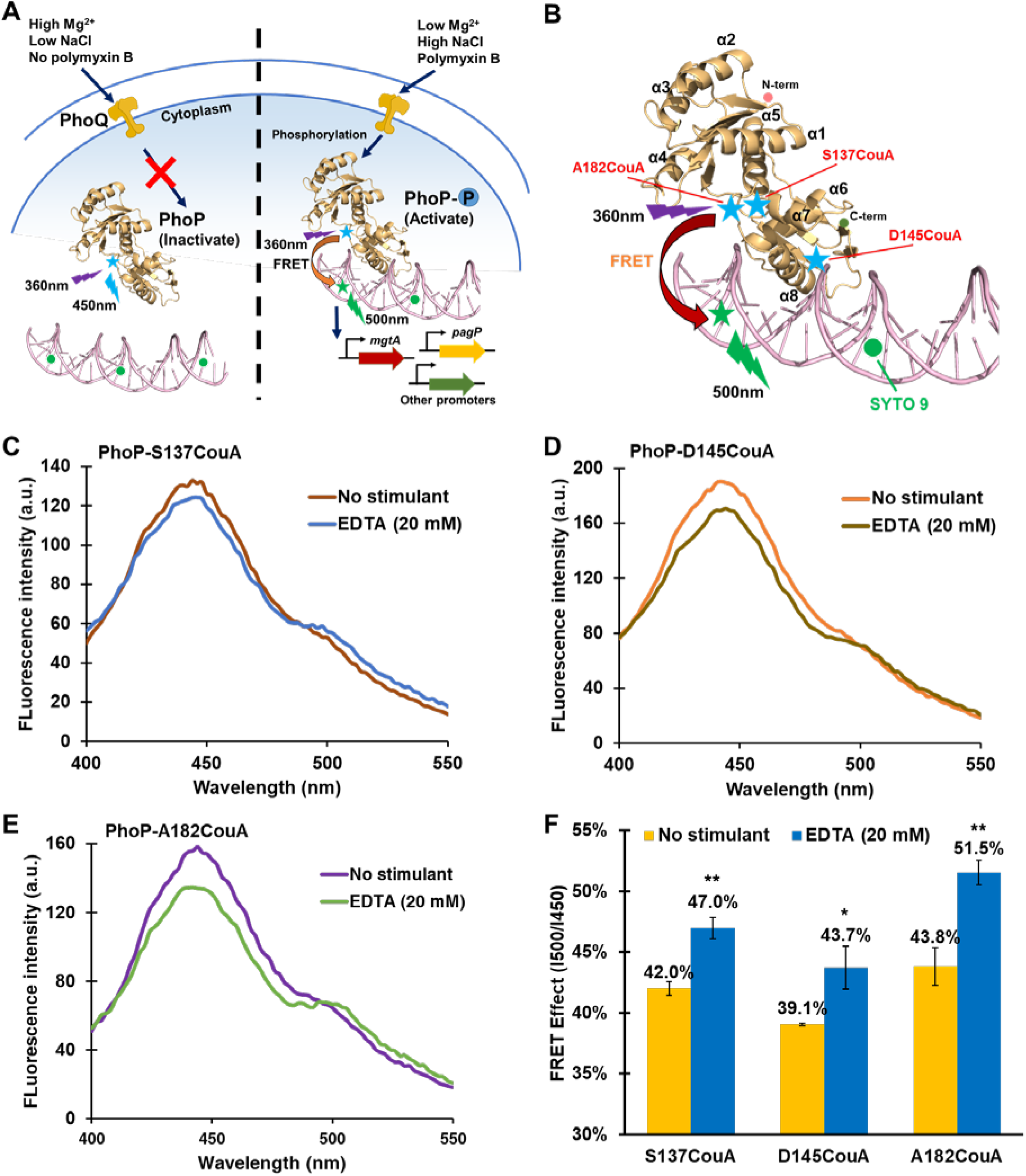
The intermolecular FRET-based TF-promoter binding assay monitors the activities of the PhoPQ TCS. (A) Diagram of the intermolecular FRET-based PhoP-promoter binding assay. Blue stars, CouA (Ex: 360 nm, Em: 450 nm); Green spheres, SYTO 9 (Ex: 483 nm, Em: 503 nm). (B) Diagram showing the locations of CouA incorporation sites in PhoP. PhoP is shown in its active state. Blue stars represent CouA, and green spheres represent SYTO 9. Ribbon structure of PhoP was simulated using the I-TASSER server. DNA fragment was adopted from Crystal structure of *Mycobacterium tuberculosis* PhoP-DNA complex (PDB ID: 5ED4). (C-E) Fluorescence spectra of BL21 (DE3) cells expressing PhoP-S137CouA (C), or PhoP-D145CouA (D), or PhoP-A182CouA (E) and stained with SYTO 9 in the presence or absence of 20 mM EDTA. (F) FRET effect of the CouA-SYTO 9 pair in the corresponding assay mixtures. Data are the mean of three biological repeats and are expressed as mean ± SD. *, *P*<0.05; **, *P*<0.01; ***, *P*<0.001, N.S., not significant relative to no EDTA treatment (based on Student’s *t* test). See also Figure S5 & S8.

Next, we employed the assay to detect the activation of the PhoPQ system by the cationic antimicrobial agent polymyxin B. Herein, the stimulant was applied to the cell suspension with a physiological level of Mg^2+^ (1 mM). We monitored time course of the FRET effected by PhoP-A182CouA and target promoters bound by SYTO 9 following applying varying concentrations of polymyxin B. Dose dependent FRET enhancement was observed at all three time courses: 10, 20, and 30 min (Fig 6A & C). The FRET enhancement maintained during the entire signal recording process (30 min), suggesting a strong and continuous activation of the PhoPQ TCS by polymyxin B. We also monitored the FRET occurrence effected by EDTA treatment in the same cell suspension, i.e., physiological level of Mg^2+^, and observed a similar pattern for FRET occurrence, degree, and dynamics (Fig 6 B & D).This result suggested that PhoPQ exerts similar signal perception and regulatory response in response to these two types of stimulants and polymyxin B activates the PhoPQ system by competing the Mg^2+^ binding site in the sensor domain of PhoP as proposed by Baber et al. (Bader et al., 2005).

**Figure 6.**
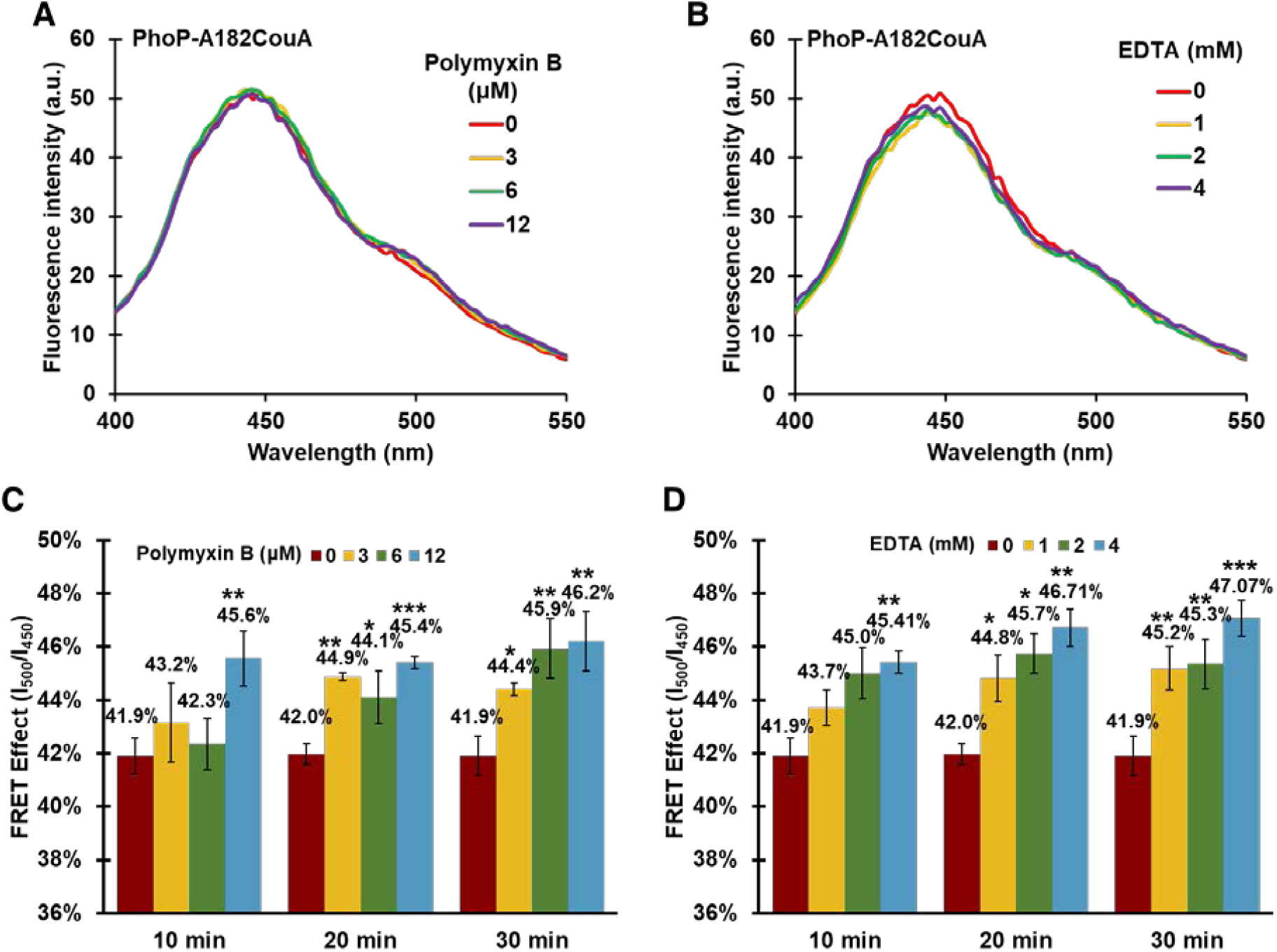
Comparison of PhoP activities in response to Mg^2+^ depletion and polymyxin B treatment. (A & B) Fluorescence spectra of BL21 (DE3) cells expressing PhoP-A182CouA (AY3709) treated by polymyxin B (A), or EDTA (B). (C & D) FRET effect of the CouA-SYTO 9 pair in the designated assay mixtures. Data are the mean of three biological repeats and are expressed as mean ± SD. *, *P*<0.05; **, *P*<0.01; ***, *P*<0.001 (based on Student’s *t* test).

### Whole-cell FRET-based TF-promoter binding assay enables identification of novel modulators of the PhoPQ TCS

The development of genome assembly, genetics, and high throughput gene expression analysis has allowed identification of a large number of TCS systems in bacterial genomes (56–58). However, a major bottleneck in understanding these signal transduction systems is the lack of information on the environmental signals and physiological conditions that trigger their activation (59, 60). In addition to reporting the in situ TF-promoter interaction in response to know stimulants of a signal transduction system, the FRET-based in vivo TF-promoter binding assay we developed can be employed to identify modulators of a regulatory system as demonstrated in the case of CueR above. To further prove this concept, we explored novel signals that modulate the PhoPQ TCS. Several previous DNA microarray studies indicated that a class of indole and its derivatives, such as indole-3-acetic acid (IAA) and tryptophan, and the aminoglycoside antibiotic gentamycin altered expression of PhoP in *E. coli* cells (61–64). It was indicated that treatment with IAA, gentamycin, and tryptophan increased the expression of PhoP, whereas indole treatment reduced its expression. To verify whether these signals indeed modulate the PhoPQ system, we treated *E. coli* cells expressing PhoP-A182CouA with these chemicals and monitored the FRET signal. As shown in Figure 7A & C, treatment with 12 mM tryptophan resulted in an enhancement of FRET (46% to 58%). Indole treatment, on the other hand, caused a slight reduction of FRET (Fig. 7B & D). Growth curve analysis indicated that 2.5 mM or above indole caused a significant reduction on the growth rate of *E. coli* (Fig. S6), suggesting that the reduction on FRET upon indole treatment may be due to perturbation on the growth of *E. coli* cells. In contrast, IAA and gentamycin treatment did not effect a FRET effect despite a broad range of concentrations and treatment time were tested (Fig. S7). These studies demonstrated that the FRET-based assay we developed is applicable to identify and verify modulators of TCSs.

**Figure 7.**
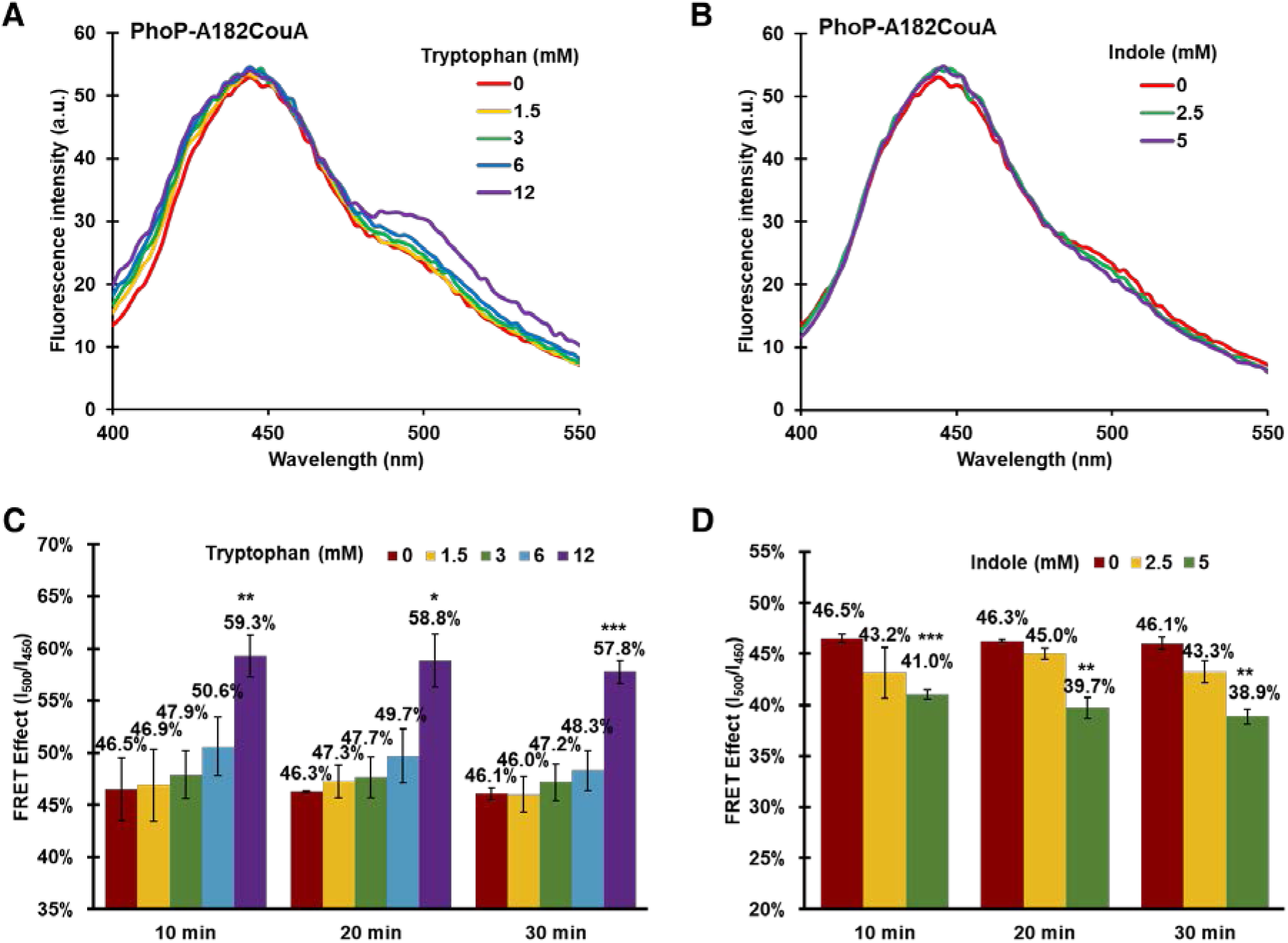
Intermolecular FRET-based assay system identifies new modulators of the PhoPQ system. (A-B) Fluorescence spectra of BL21 (DE3) cells expressing PhoP-A182CouA (AY3709) in the presence of tryptophan (A), or indole (B). (C-D) FRET effect of assay mixtures under the designated conditions. Data are the mean of three biological repeats and are expressed as mean ± SD. *, *P*<0.05; **, *P*<0.01; ***, *P*<0.001 (based on Student’s *t* test). See also Figure S6 & S7.

### FRET-based TF-promoter assay monitors the activation of the PhoPQ system in E. coli cells colonized in the C. elegans gut

Next, we expanded the assay system to monitor the activation of a signal transduction system and TF activities in bacteria colonized in an animal host using *C. elegans* as a paradigm. *C. elegans* is a model organism widely employed to study bacteria-host interaction due to its invariant cell lineage, simple body structure, and short life cycle (65, 66). The transparent body of the *C. elegans* is particularly ideal for spectroscopic analysis. Since PhoPQ serves as the primary regulatory system during the adaptation of enterobacteria to their host environments (67–69), we tested the application of the FRET-based assay to report the PhoP-promoters interaction in *E. coli* colonized in the *C. elegans* gut. We fed the *C. elegans* L4 stage larva with *E. coli* BL21 cells expressing PhoP-A182CouA (AY3706) and stained with SYTO 9 (Fig. 8A)*. C. elegans phm-2*(-) mutant animals (DA597) were used in the experiment because they have a dysfunctional grinder (70), allowing the observation of intact *E. coli* in *C. elegans* gut. *C. elegans* fed with the *E. coli* OP50 served as a control (Fig. 8B). The expected blue fluorescence (420 to 480 nm) (Fig. 8C, upper left panel) and green fluorescence (490 to 600nm) (Fig. 8C, upper right panel) which corresponds to the emission of CouA and SYTO 9, respectively, was observed in the *C. elegans* intestinal tract when the animals were excited at 405 nm and 488 nm, respectively. Particularly, the fluorescence in the gut was shown as small puncta rather than being diffused, indicating the presence of intact *E. coli* cells emitting fluorescence of PhoP-A182CouA and SYTO 9-bound with DNA in the *C. elegans* gut.

**Figure 8.**
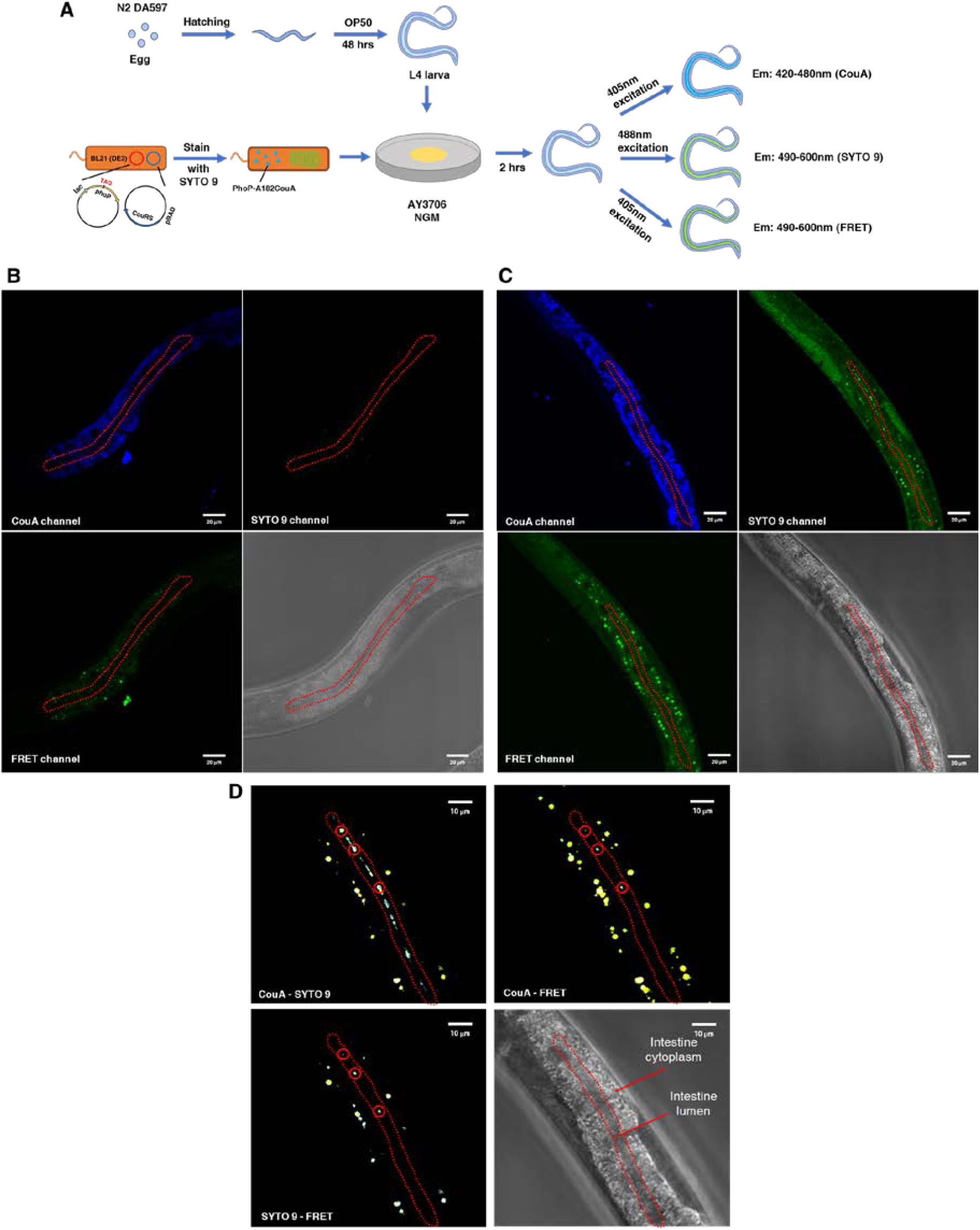
FRET measurement reports the PhoP-promoter activitiy in *E. coli* cells colonized in the *C. elegans* gut. (A) Workflow of the intermolecular FRET-based assay in the *C. elegans* colonization model. (B & C) Fluorescence of *C. elegans* fed with OP50 (B) or BL21 (DE3) cells expressing PhoP-A182CouA (AY3709) (C) examined by confocal fluorescent microscope. Upper left: CouA emission fluorescence recorded in the range of 420-480 nm upon excitation at 405 nm. Upper right: SYTO 9 emission fluorescence recorded in the range of 490-600 nm upon excitation at 488 nm. Lower left: FRET signal recorded in the range of 490-600 nm upon excitation at 405 nm. Lower right: Differential interference contrast (DIC) image of the *C. elegans* intestine area subjected to fluorescence recording. (D) Colocalization of different fluorescent channels in the intestine area of *C. elegans* in (C). Red circles indicate the colocalization between the indicated fluorescent channels. Upper left: CouA channel and SYTO 9 channel. Upper right: CouA channel and FRET channel. Lower left: SYTO 9 channel and FRET channel. Lower right: DIC image showing the intestine lumen and cytoplasm area. Red dash circles represent the intestine lumen region. See also Figure S5.

To examine whether PhoPQ system is activated to bind its target promoters and regulate gene expression in the *C. elegans* gut, we excited the *C. elegans* at 405 nm and recorded the green fluorescence emitted by SYTO 9-bound with DNA in the *C. elegans* intestinal lumen (490 to 600 nm) (Fig. 8C, lower left panel), i.e., FRET effect. FRET signals were recorded in several focal points in the gut lumen, indicating the binding of PhoP-A182CouA with SYTO 9 stained targeting promoters in these *E. coli* cells (Fig. 8C, lower left panel). Colocalization of the signals corresponding to the emission fluorescence of CouA and SYTO 9 with that of FRET further confirmed this result (Fig. 8D). Hence, the method we developed is applicable to examine activities of transcription factors at the bacteria-host interface.

## Discussion

In recent decades, genetic codon expansion technique, which enables site-specific incorporation of genetically encoded unnatural amino acids (UAA) into proteins, has revolutionized our ability to investigate the functions of proteins and their dynamics in living systems (71–74). However, the technique has not been leveraged to study transcription regulation during the stress adaptation in bacteria which is essential to bacterial survival and infection. In this study, we encoded a bright fluorescent unnatural amino acid (FUAA) CouA site-specifically into the DNA-binding domain of bacterial transcription regulator proteins and monitored the TF-promoter interaction in response to various stimulants and stresses based on the intermolecular FRET between the FUAA and the live cell DNA stain SYTO 9. As FRET represents almost the most accessible, sensitive, and super-resolved measurement for macromolecule associations, the assay system provides a powerful tool to monitor the adaptive gene regulation in living bacteria. We showed that the intermolecular FRET-based assay is capable of monitoring the binding of TF with its target promoters and activation of the corresponding regulatory systems in diverse signal transduction processes with specificity and sensitivity. Furthermore, the assay system is applicable to screen modulators of the regulatory systems of interests and report the activities of transcription factors in bacteria colonized in the *C. elegans* host. Hence, it provides a robust TF-promoter binding assay for investigating the complex and dynamic signal transduction processes in living cells which are inaccessible by other methods currently available.

A common concern about intermolecular FRET-based biosensors is the occurrence of background noise and false positives caused by over-production of the donor and acceptor fusions and cross-excitation of the acceptor by the excitation light of the donor. The problem of cross-excitation can be effectively solved by selecting a donor fluorophore with a large Stokes shift (75). The FRET donor we employed in this study, CouA, is a fluorophore with characteristic features ideal for FRET owing to its small size, high quantum field, and large Stokes shift (27). Indeed, no cross-excitation occurred in both the in vitro and in vivo FRET measurements established herein as demonstrated by a lack of fluorescence signal at 500 nm in the mixture of the CueR-F58CouA protein with a series concentration of SYTO 9 (Fig. S3A) and a modest increase of fluorescence intensity at 500 nm in the whole-cell suspension of BL21 mixed with varying amounts of SYTO 9 (Fig. S3B). A moderate condition for protein expression and CouA incorporation was applied for the whole-cell FRET-based assay. The level of CouA-incorporated TF proteins in the cell was determined to be in the range of 100 μg/ml which affords sufficient fluorescence for FRET measurements but retains ample sensitivity as demonstrated by the assay of three different regulatory systems CueR, BasSR, PhoPQ in response to a broad range of signals and stresses tested. SYTO 9 is a common nucleic acids stain widely applied to identify and enumerate live bacteria in a population. Conceivably, the DNA content of a cell influences and determines the amount of SYTO 9 required to sufficiently cover the targeting promoters of the TF protein of interest. However, studies from McGoverin et al. (42) showed that nucleobase length and GC content of a bacterium did not significantly affect the SYTO 9 concentration required for enumeration of bacteria. It was proposed that the saturation concentration of SYTO 9 for a specific bacterial species was largely determined by the permeability of SYTO 9 to the bacterial cells and that micromolar (1 μM) SYTO 9 affords a robust signal for enumeration of common bacterial species including *E. coli* (42). In agreement with this notion, a limited extent of fluorescence intensity change was observed in the *E. coli* BL21 whole-cell suspension when 0.5-20 μM SYTO 9 was supplied. The SYTO 9 concentration we selected (7.5 μM) afforded sufficiently bright fluorescence for both fluorimeter and fluorescent microscopy analysis and retained ample sensitivity to report the FRET occurrence and its degree in response to stress signals. Hence, the intermolecular FRET-based assay we developed provides an accessible, robust, and generalizable approach to monitor TF-promoter interactions in living bacteria.

In most applications involving the site-specific incorporation of UAA for functional characterization of proteins, the 3D structure of the proteins is often known and is required. Using the CueR transcription factor as an example, we demonstrated that structure-guided determination of CouA incorporation sites facilitated the establishment of the intermolecular FRET-based TF-promoter binding assay. However, crystal structures of many important transcription regulators and those of newly annotated regulators are unknown. Using BasR and PhoP as examples, we showed that simulated structures are reliable and applicable for determining the CouA incorporation sites for the assay. In these applications, 3D structures of BasR and PhoP were simulated by I-TASSER using the crystal structure of the corresponding homologue proteins, i.e., *K. pneumoniae* PmrA and *M. tuberculosis* PhoP, respectively, as the template. Our further analysis showed that protein 3D structures predicted by the AI-based programs such as AlphaFold and tFold (76, 77) in the absence of an existing template are also reliable and applicable for determination of the CouA incorporation sites (Fig. S8). Our studies demonstrated that the loop regions between α helices or β sheets in the DBD are optimal locations for CouA incorporation to monitor the TF-promoter interaction. However, certain amino acid residues, such as proline, is not tolerant to replacement by UAAs. Despite this, the fact that CouA can be incorporated into multiple locations in the loop regions of DBD to report the TF-promoter binding in CueR, BasR and PhoP demonstrated the robustness and accessibility of the assay system we developed.

Two major types of signal transduction systems existed in bacteria, the cytoplasmic one-component transcription regulators which primarily sense intracellular signals and the membrane-associated two-component systems which primarily sense membrane, periplasmic, and extracellular signals and stimulants. We showed that supplying inducing signals or stimulants in the assay suspension expressing the CouA-incorporated RR of TCSs elicited instant FRET. However, when the whole-cell assay was employed to monitor the binding of cytoplasmic TF CueR with its target promoters, supplementing the stimulant in the assay suspension did not yield expected FRET signal even after 40 min incubation (data not shown). Instead, when the stimulants or modulators were supplemented during CouA incorporation and protein expression, distinct FRET effect (in the presence of inducing signals) or abolishment of FRET (in the presence of signals inactivating the regulator) was observed. Hence, the stimulant application step needs to be adjusted depending on the types of the regulatory systems being investigated to enable proper signal perception.

In the past decades, the development of synthetic strategies has greatly accelerated the design and preparation of new FUAAs and nucleic acid dyes (78). With the development of these agents that display better fluorescence features, environmental sensitivity, and cell compatibility, the FRET-based assay for TF-promoter binding, and to a larger extent of protein-DNA interaction, will be continuously optimized for sensitive, robust, and broad applications.

## Data availability

The mass spectrometry proteomics data have been deposited to the ProteomeXchange Consortium via the PRIDE(79) partner repository with the dataset identifier PXD030973. (Username, Password: OV7ZS40g)

## Supporting information

Protein-DNA models

## Acknowledgement

The *C. elegans* strains used in this study were provided by the Caenorhabditis Genetics Center, which is funded by NIH Office of Research Infrastructure Programs (P40 OD010440). We thank to Dr. Chenghao Bi (Tianjin Institute of Industrial Biotechnology, Chinese Academy of Sciences) for the gift of the pCAGO plasmid. We thank to Proteomics and Metabolomics Core, Centre for PanorOmic Sciences in HKU for their help and support on proteomics analysis.

