## Supplemental figures and tables for "FRET monitoring of transcription factor activities in living bacteria"

**Supplementary data**

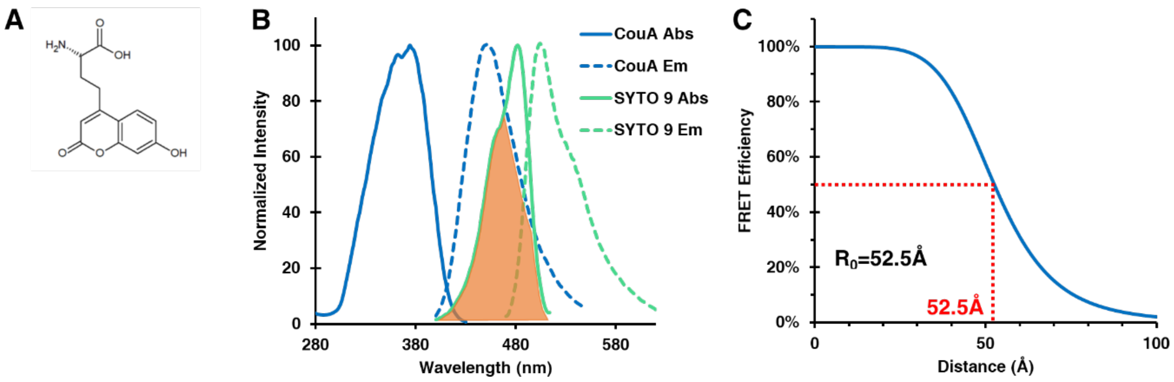


Figure S1. Fluorescence properties of the FRET pair employed in this study, CouA and SYTO 9. (A) Chemical structure of L-(7-hydroxylcoumarin-4-yl) ethylglycine (CouA). (B) Normalized fluorescence spectra of CouA (blue) and SYTO 9. (C) Graph showing the relationship between FRET efficiency and the distance between CouA and SYTO 9. Abs, absorption; Em, emission.

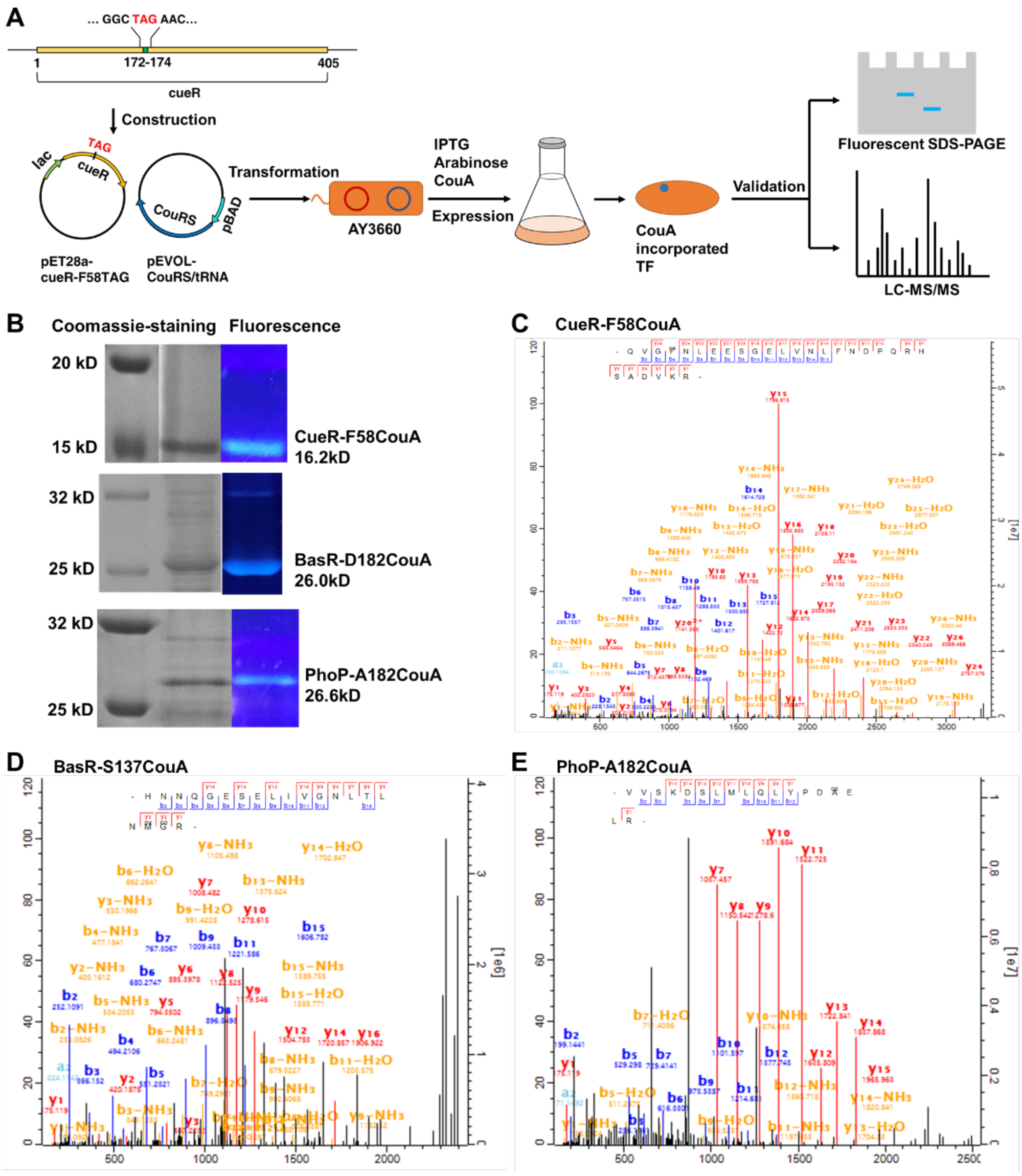


Figure S2. Expression and verification of CouA incorporation into TF proteins. (A) Diagram of the workflow for CouA incorporation into TF proteins employing the genetic code expansion strategy. (B) SDS-PAGE analysis of the designated proteins visualized by Coomassie blue stain and under UV light. (C-E) LC MS/MS spectra of CueR-F58CouA (C), BasR-S137CouA (D), and PhoP-A182CouA (E). B series show the masses of peptides in the forward amino acid sequence, y series show the masses of peptides in the reverse amino acid sequence.

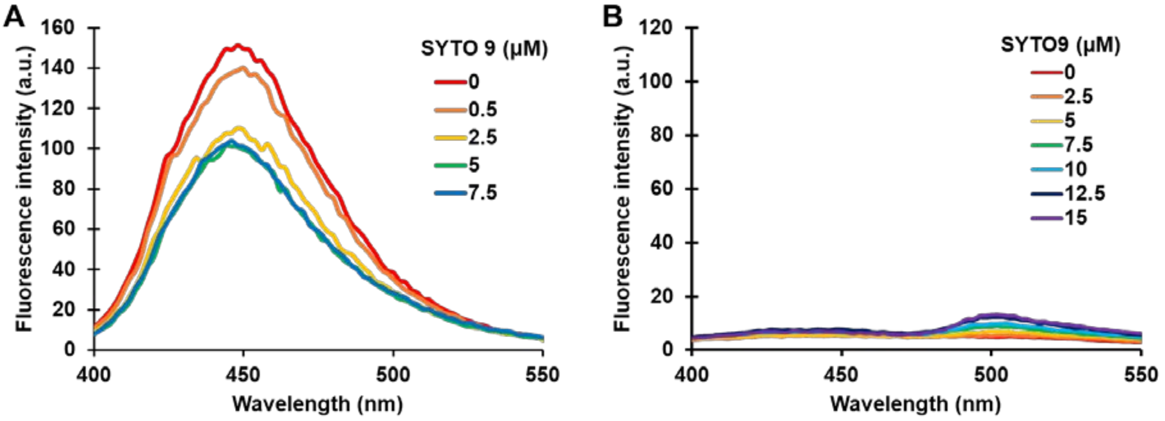


Figure S3. (A) Fluorescence spectra of purified CueR-F58CouA protein (10 μM) mixed with varying amount of SYTO 9 (0-7.5 μM). (B) Fluorescence spectra of *E. coli* BL21 cells without induction and stained with varying amount of SYTO 9 (0-15 μM) excited by 360 nm light source.

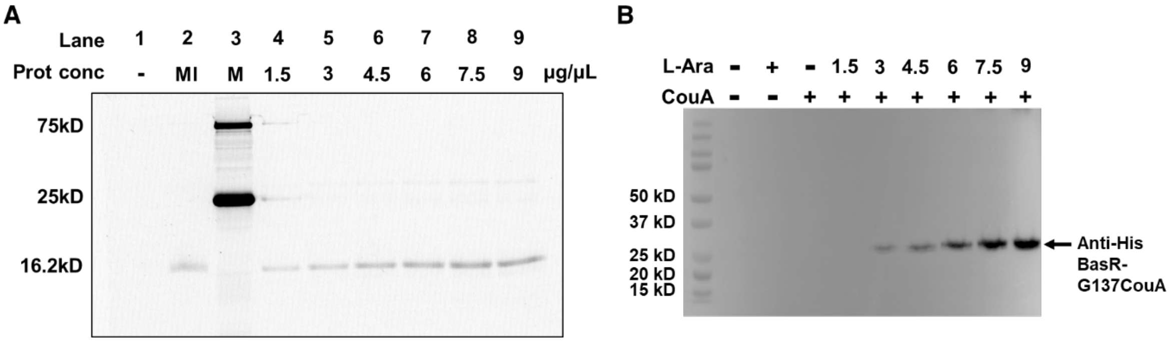


Figure S4. TF protein levels in the whole-cell intermolecular FRET assay. (A) Fluorescent SDS-PAGE image of the CueR-F58CouA protein. Lane 1, negative control; Lane 2, cell lysate of AY3660 cells mildly induced by supplementing 0.5 mM IPTG, 0.4% L-arabinose and 1 mM CouA at 37°C for 4 hrs; Lane 3, protein marker; Lane 4-9, purified CueR-F58CouA with designated concentrations. (B) Western blot image of BasR-G137CouA. Lane 1, protein marker; Lane 2-4, negative controls; Lane 5-10, cell lysates of AY7065 cells subjected to induction by supplementing a series concentrations of L-arabinose in the presence of 1 mM CouA at 37°C for 5 hrs.

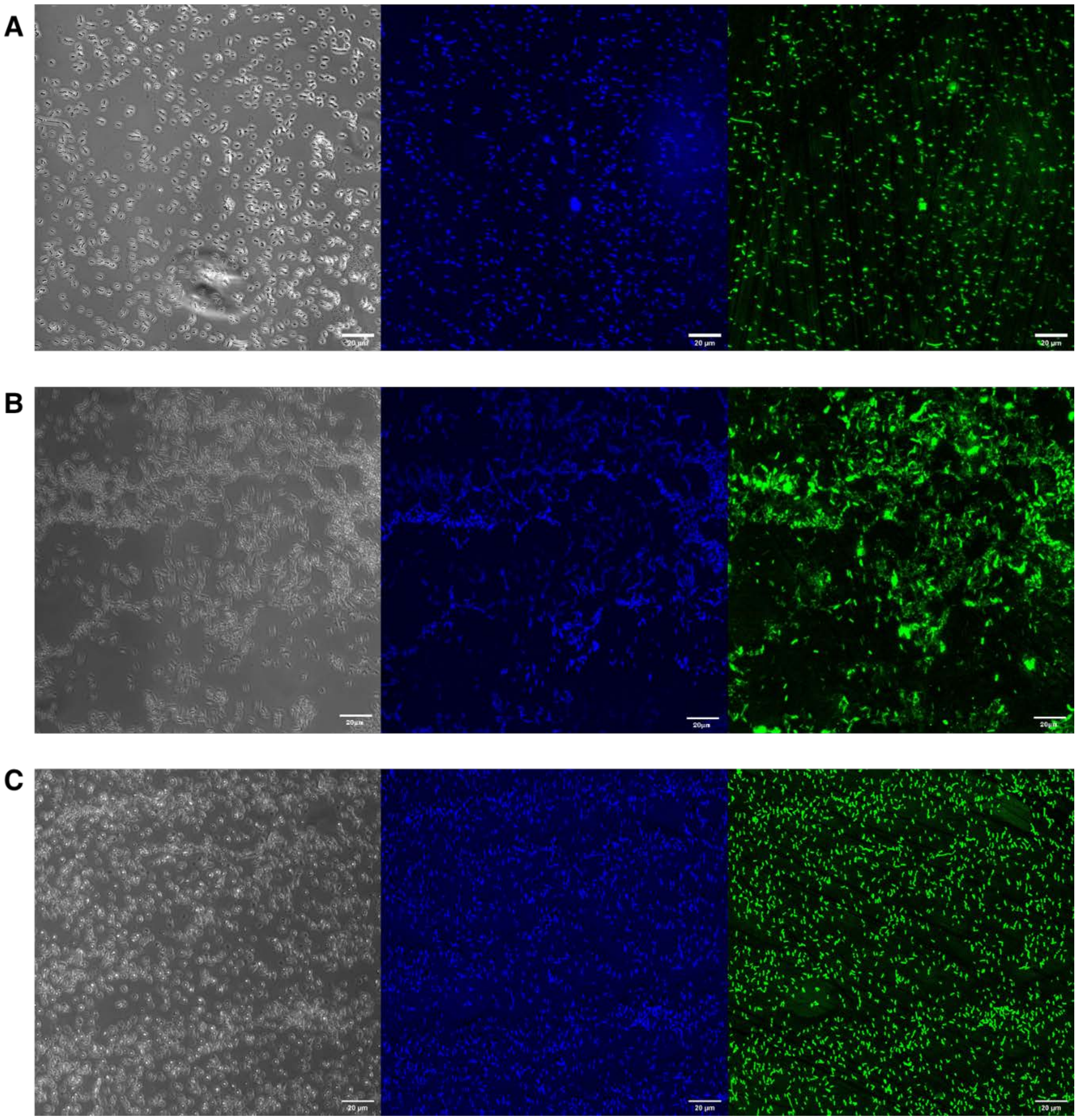


Figure S5. Fluorescence images of *E. coli* BL21(DE3) or MG1655 cells expressing CouA incorporated TF proteins (blue) and stained with SYTO 9 recorded by confocal microscope. Left, middle and right images represent that obtained in DIC, CouA, SYTO 9 emission channels, respectively. (A) BL21 cells expressing CueR-F58CouA. (C) MG1655 cells expressing BasR-D182CouA. (B) BL21 cells expressing PhoP-A182CouA.

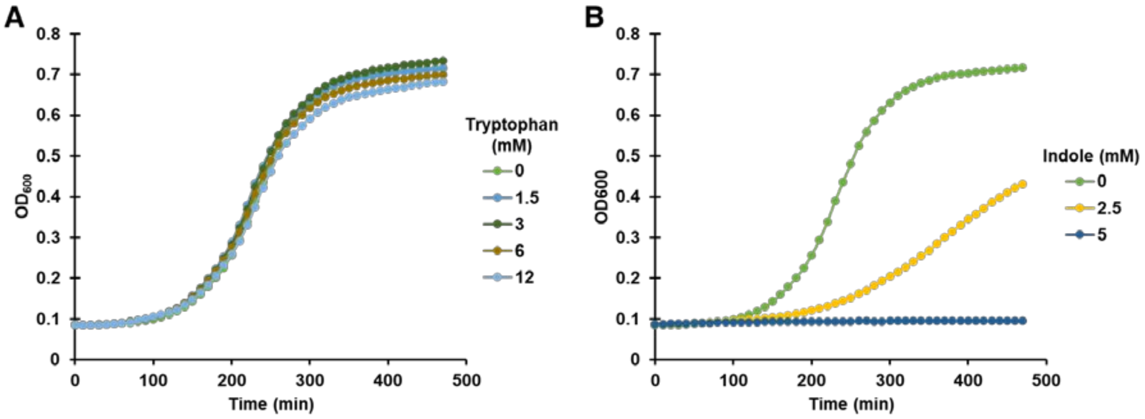


Figure S6. Growth curves of *E. coli* AY3709 under different concentration of tryptophan (A) or indole (B).

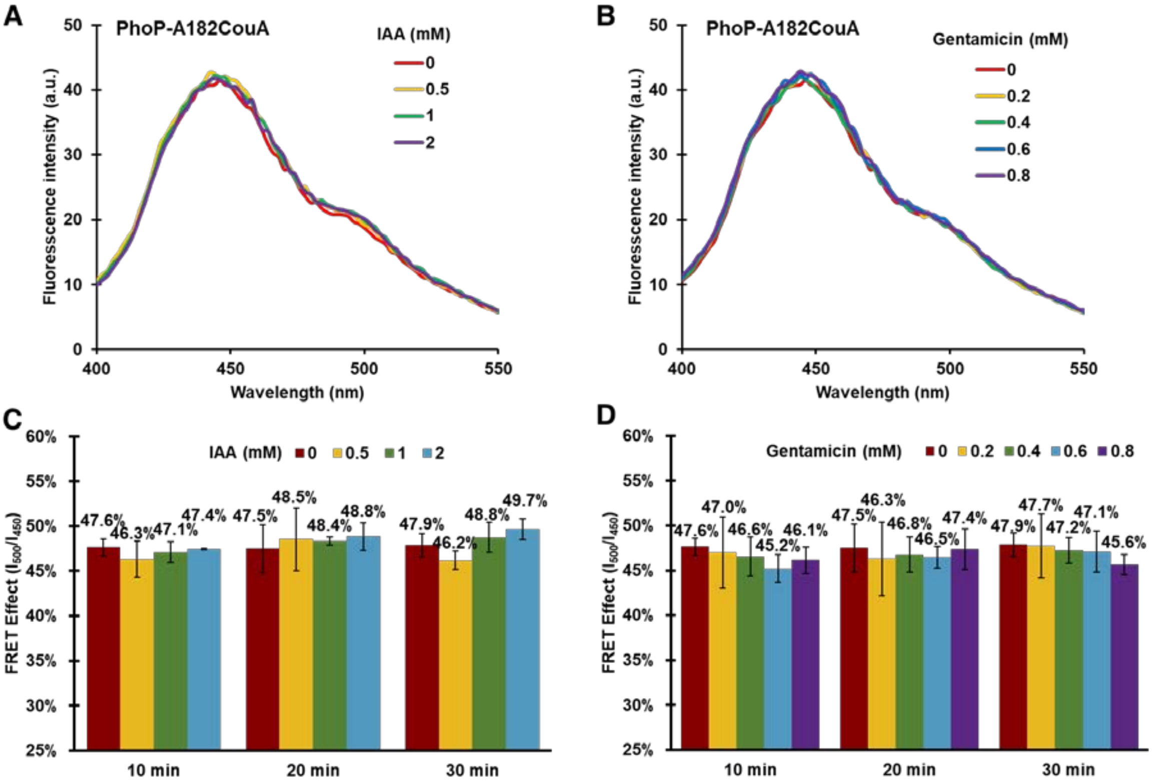


Figure S7. Intermolecular FRET-based assay indicated that IAA and gentamycin did not modulate activity of the PhoPQ system. (A-B) Fluorescence spectra of BL21 (DE3) cells expressing PhoP-A182CouA (AY3709) in the presence of IAA (0, 0.5, 1, 2 mM) (A), or gentamicin (0, 0.2, 0.4, 0.6, 0.8 mM) (B). (C-D) FRET effect of CouA-SYTO 9 in BL21 (DE3) cells expressing PhoP-A182CouA (AY3709) under the designated conditions. Data are the mean of three biological repeats and are expressed as mean ± SD.

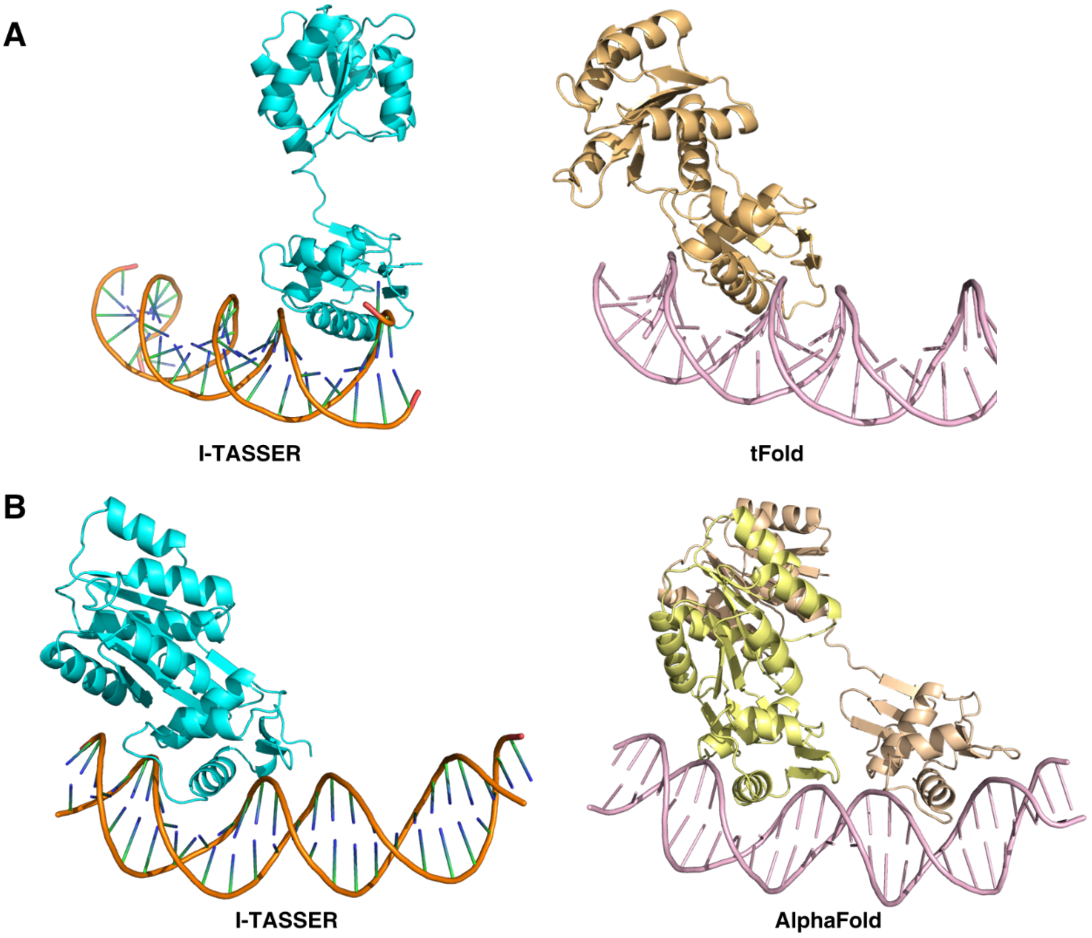


C

| Incorporation site in PhoP | Distance in I-TASSER (Å) | Distance in tFold (Å) |
|---|---|---|
| S137 | 25.1 | 26.4 |
| D145 | 30.0 | 32.4 |
| A182 | 20.2 | 25.3 |

| Incorporation site in BasR | Distance in I-TASSER (Å) | Distance in AlphaFold (Å) |
|---|---|---|
| G137 | 26.3 | 31.1 |
| E146 | 28.1 | 27.6 |
| S167 | 23.8 | 25.5 |
| D182 | 21.3 | 19.6 |

Figure S8. Comparison of the 3D structures of *E. coli* PhoP and BasR proteins predicted by tFold and AlphaFold with those simulated by I-TASSER. (A) Predicted 3D structure of PhoP. Left, simulated by I-TASSER; Right, predicted by tFold. (B) Predicted 3D structure of BasR. Left, simulated by I-TASSER; Right, predicted by AlphaFold. (C) The calculated distance of CouA incorporation sites to the closest DNA minor groove in the designated 3D structure predictions.

**Table S1.** Properties of fluorescent unnatural amino acids with available aaRS-tRNA pairs

| UAA | Formal name | $\lambda_{abs}$ (nm) | $\lambda_{em}$ (nm) | ϕ | ε ($cm^{-1}M^{-1}$) |
|---|---|---|---|---|---|
| TerphenylA | 4-biphenyl-l-phenylalanine | 280 | 342 | 0.49 | - |
| CouA | l-(7-hydroxycoumarin-4-yl)ethylglycine | 360 | 450 | 0.63 | 17000 |
| Lys-CouA | Coumarin lysine analogues | 360 | 450 | - | - |
| ACD | acridon-2-ylalanine | 386 | 446 | 0.95 | |
| ANAP | 3-(6-acetylnaphthalen-2-ylamino)-2-aminopropanoic acid | 360 | 490 | 0.48 | 17500 |
| Dansylalanine | amino-3-(5-(dimethylamino)naphthalene-1-sulfonamide)propanoic acid | 386 | 540 | - | - |

$\lambda_{abs}$, absorption wavelength, $\lambda_{em}$, emission wavelength, ϕ, quantum yield, ε, extinct coefficient.

**Table S2.** Primers used in this study

| Primer | DNA sequences |
|---|---|
| cueR-F | 5'-CATGCCATGGCCATGAACATCAGCGATGTAGC-3' |
| cueR-R | 5'-CCGCTCGAGCCCTGCCCGATGATGACAGC-3' |
| cueR-F58TAG-F | 5'-GCAGGTGGGCTAGAACCTGGAAGAGAGC-3' |
| cueR-F58TAG-R | 5'-CTTCCAGGTTCTAGCCCACCTGCCGTGC-3' |
| cueR-T27TAG-F | 5'-GGGGCTGGTGTAGCCGCCGATGCGCAGC-3' |
| cueR-T27TAG-R | 5'-GCATCGGCGGCTACACCAGCCCCTTCTC-3' |
| C-Y39-F | 5'-ACGAACTGACCTTACTGCGCCAGGC-3' |
| C-Y39-R | 5'-GATAACCGTTTTCGCTGCGCATCGG-3' |
| L-Y39-F | 5'-GCGCAGCGAAAACGGTTATCGCACCTAGACGCAGCAGCATCTCAACGAACTGACCTTAC-3' |
| L-Y39-R | 5'-GTAAGGTCAGTTCGTTGAGATGCTGCTGCGTCTAGGTGCGATAACCGTTTTCGCTGCGC-3' |
| C-Q74-F | 5'-CGTCAAACGGCGCACGCTGG-3' |
| C-Q74-R | 5'-ATTCACCAGCTCGCCGCTCT-3' |
| L-Q74-F | 5'-GGCGAGCTGGTGAATCTGTTTAACGACCCGTAGCGGCACAGCGCCGACGTCAAACGGCG-3' |
| L-Q74-R | 5'-CGCCGTTTGACGTCGGCGCTGTGCCGCTACGGGTCGTTAAACAGATTCACCAGCTCGCC-3' |
| pcopA-F | 5'-CCCGCAACTTAACTACAGG-3' |
| pcopA-R | 5'-GTGACATAAAACACTCCTTTAAG-3' |
| Frag1-F | 5'-AACATCAGCGATGTAGCAAA-3' |
| Frag1-R | 5'-TCAGTTCGTTGAGATGCTGC-3' |
| cueR-R18A-F | 5'-CAAAGCCATTGCCTTCTATGAAGAGAAG-3' |
| cueR-R18A-R | 5'-CTTCATAGAAGGCAATGGCTTTGCTGGT-3' |
| cueR-R37A-F | 5'-AAACGGTTATGCCACCTACACGCAGCAG-3' |
| cueR-R37A-R | 5'-GCGTGTAGGTGGCATAACCGTTTTCGCT-3' |
| phoP-F | 5'-CATGCCATGGCCATGCGCGTACTGGTTGTTGA-3' |
| phoP-R | 5'-CCGCTCGAGGCGCAATTCGAACAGATAGC-3' |
| phoP-A182TAG-F | 5'-CTATCCGGATTAGGAGCTGCGGGAAAGC-3' |
| phoP-A182TAG-R | 5'-CCCGCAGCTCCTAATCCGGATAGAGTTG-3' |

| | |
|---|---|
| phoP-S137TAG-F | 5'-GGTTGATCTCTAGCGCCGTGAATTATCT-3' |
| phoP-S137TAG-R | 5'-ATTCACGGCGCTAGAGATCAACCTGAAA-3' |
| phoP-D145TAG-F | 5'-ATCTATTAATTAGGAAGTGATCAAACTG-3' |
| phoP-D145TAG-R | 5'-TGATCACTTCCTAATTAATAGATAATTC-3' |
| basR-F | 5'-AGAAGGAGATATACCATGGCCATGAAAATTCTGATTGTTGAA-3' |
| basR-R | 5'-GGTGGTGGTGCTCGAGGTTTTCCTCATTCGCGACCA-3' |
| basR-G137TAG-F | 5'-GCTGAACATGTAGCGCCGTCAGGTATGG-3' |
| basR-G137TAG-R | 5'-CCTGACGGCGCTACATGTTCAGCGTCAG-3' |
| basR-E146TAG-F | 5'-GATGGGCGGTTAGGAGTTGATTCTGACG-3' |
| basR-E146TAG-R | 5'-GAATCAACTCCTAACCGCCCATCCATAC-3' |
| basR-S167TAG-F | 5'-CAAAGCAGGCTAGCCGGTGCATCGGGAA-3' |
| basR-S167TAG-R | 5'-GATGCACCGGCTAGCCTGCTTTGAGCAT-3' |
| basR-D182TAG-F | 5'-CTATAACTGGTAGAATGAACCCTCGACC-3' |
| basR-D182TAG-R | 5'-AGGGTTCATTCTACCAGTTATAGATGTC-3' |

**Table S3.** Bacterial strains and plasmids used in this study

| **Strain/Plasmid** | **Genotype** | **Source** |
|---|---|---|
| MG1655 | *E. coli F- λ- ilvG- rfb-50 rph-1* | Lab collection |
| DH5α | *E. coli F*$^{–}$ *endA1 glnV44 thi-1 recA1 relA1 gyrA96 deoR nupG purB20 φ80dlacZΔM15 Δ(lacZYA-argF)U169, hsdR17*($r_K^{–}m_K^{+}$), *λ*$^{–}$ | Lab collection |
| BL21(DE3) | *E. coli* B F$^{–}$ *ompT gal dcm lon hsdS*$_B$($r_B^{–}m_B^{–}$) [*malB*$^{+}$]$_{K\text{-}12}$($λ^S$) | Lab collection |
| OP50 | *E. coli ura-, strR, rnc-, (delta)attB::FRT-lacI-lacUV5p-T7* | Lab collection |
| AY3700 | *E. coli* DH5α, pET28a-*cueR* | This study |
| AY3701 | *E. coli* DH5α, pET28a-*cueR*-F58TAG | This study |
| AY3702 | *E. coli* DH5α, pEVOL-CouRS | Lab collection |
| AY3619 | *E. coli* BL21, pET28a-*cueR* | This study |
| AY3660 | *E. coli* BL21, pET28a-*cueR*-F58TAG and pEVOL-CouRS | This study |
| AY3703 | *E. coli* DH5α, pET28a-*phoP* | This study |
| AY3704 | *E. coli* DH5α, pET28a-*phoP*-S137TAG | This study |
| AY3705 | *E. coli* DH5α, pET28a- *phoP*-D145TAG | This study |
| AY3706 | *E. coli* DH5α, pET28a- *phoP*-A182TAG | This study |
| AY3707 | *E. coli* BL21, pET28a- *phoP*-S137TAG and pEVOL-CouRS | This study |
| AY3708 | *E. coli* BL21, pET28a- *phoP*-D145TAG and pEVOL-CouRS | This study |
| AY3709 | *E. coli* BL21, pET28a- *phoP*-A182TAG and pEVOL-CouRS | This study |
| AY3710 | *E. coli* BL21, pET28a-*phoP* | This study |
| AY3711 | *E. coli* DH5α, pET28a-*cueR*-T27TAG | This study |
| AY3712 | *E. coli* DH5α, pET28a-*cueR*-Y39TAG | This study |
| AY3713 | *E. coli* DH5α, pET28a-*cueR*-Q74TAG | This study |
| AY3714 | *E. coli* BL21, pET28a-*cueR*-T27TAG and pEVOL-CouRS | This study |
| AY3715 | *E. coli* BL21, pET28a-*cueR*-Y39TAG and pEVOL-CouRS | This study |
| AY3716 | *E. coli* BL21, pET28a-*cueR*-Q74TAG and pEVOL-CouRS | This study |
| AY3717 | *E. coli* DH5α, pET28a-*cueR*-F58TAG-R18A | This study |
| AY3718 | *E. coli* DH5α, pET28a-*cueR*-F58TAG-R37A | This study |
| AY3719 | *E. coli* BL21, pET28a-*cueR*-F58TAG-R18A and pEVOL-CouRS | This study |
| AY3720 | *E. coli* BL21, pET28a-*cueR*-F58TAG-R37A and pEVOL-CouRS | This study |
| AY3616 | *E. coli* MG1655, Pnn387-$P_{cueO}$-*lacZ* | Lab collection |
| AY4426 | *E. coli* MG1655, Pnn387-$P_{copA}$-*lacZ* | Lab collection |
| AY7030 | *E. coli* MG1655 *ΔbasR* | Lab collection |
| AY7031 | *E. coli* DH5α, pET28a-pBAD-*basR* | This study |
| AY7032 | *E. coli* DH5α, pET28a-pBAD-*basR*-D182TAG | This study |
| AY7033 | *E. coli* DH5α, pET28a-pBAD-*basR*-G137TAG | This study |
| AY7034 | *E. coli* DH5α, pET28a-pBAD-*basR*-E146TAG | This study |
| AY7035 | *E. coli* DH5α, pET28a-pBAD-*basR*-S167TAG | This study |
| AY7036 | *E. coli* DH5α, pET28a-pBAD-*basR*-3×FLAG | This study |
| AY7037 | *E. coli* BL21 $ΔP_{cueO}$ | This study |

| | | |
|---|---|---|
| AY7038 | *E. coli* BL21 $\Delta P_{cueO}$ $\Delta P_{copA}$ | This study |
| AY7063 | *E. coli* MG1655 *ΔbasR*, pET28a-pBAD-*basR* and pEVOL-CouRS | This study |
| AY7064 | *E. coli* MG1655 *ΔbasR*, pET28a-pBAD-*basR*-D182TAG and pEVOL-CouRS | This study |
| AY7065 | *E. coli* MG1655 *ΔbasR*, pET28a-pBAD-*basR*-G137TAG and pEVOL-CouRS | This study |
| AY7066 | *E. coli* MG1655 *ΔbasR*, pET28a-pBAD-*basR*-E146TAG and pEVOL-CouRS | This study |
| AY7067 | *E. coli* MG1655 *ΔbasR*, pET28a-pBAD-*basR*-S167TAG and pEVOL-CouRS | This study |
| AY7068 | *E. coli* MG1655 *ΔbasR*, pET28a-pBAD-*basR*-3×FLAG | This study |
| pAY3702 | pEVOL-CouRS | Lab collection |
| pAY3619 | pET28a-*cueR* | This study |
| pAY3703 | pET28a-*phoP* | This study |
| pAY3701 | pET28a-*cueR*-F58TAG | This study |
| pAY3704 | pET28a-*phoP*-S137TAG | This study |
| pAY3705 | pET28a- *phoP*-D145TAG | This study |
| pAY3706 | pET28a- *phoP*-A182TAG | This study |
| pAY3711 | pET28a-*cueR*-T27TAG | This study |
| pAY3712 | pET28a-*cueR*-Y39TAG | This study |
| pAY3713 | pET28a-*cueR*-Q74TAG | This study |
| pAY7031 | pET28a-pBAD-*basR* | This study |
| pAY7032 | pET28a-pBAD-*basR*-D182TAG | This study |
| pAY7033 | pET28a-pBAD-*basR*-G137TAG | This study |
| pAY7034 | pET28a-pBAD-*basR*-E146TAG | This study |
| pAY7035 | pET28a-pBAD-*basR*-S167TAG | This study |
| pAY7036 | pET28a-pBAD-*basR*-3×FLAG | This study |
